# Locus coeruleus activation ‘resets’ hippocampal event representations and separates adjacent memories

**DOI:** 10.1101/2024.08.15.608148

**Authors:** David Clewett, Ringo Huang, Lila Davachi

## Abstract

Memories reflect the ebb and flow of experiences, capturing distinct events from our lives. Using a combination of functional magnetic resonance imaging (fMRI), neuromelanin imaging, and pupillometry, we show that arousal and locus coeruleus (LC) activation segment continuous experiences into discrete memories. As sequences unfold, encountering a context shift, or event boundary, triggers pupil-linked arousal and LC processes that predict later memory separation. Boundaries furthermore promote temporal pattern separation within left hippocampal dentate gyrus, which correlates with heightened LC responses to those same transition points. Unlike transient LC effects, indirect structural and functional markers of elevated background LC activation correlate with reduced arousal-related LC and pupil responses at boundaries, suggesting that hyperarousal disrupts event segmentation. Our findings support the idea that arousal mechanisms initiate a neural and memory ‘reset’ in response to significant changes, fundamentally shaping the episodes that define episodic memory.

## Introduction

As time passes, we are exposed to a continuous stream of information. To make sense of it all, individuals tend to group similar information into distinct mental ‘episodes’ based on context, like place or time^1^. For instance, being in your kitchen helps create a memory of “breakfast” by linking together various details like eggs, the table, and time of day. Conversely, transitioning to a new context or situation, like leaving home for work, facilitates perception of a new event^1,2^. Memory formation therefore involves a delicate balance between integrating continuous information and segmenting distinct episodes in perception and memory, often triggered by context shifts that act as event boundaries^3–5^. A wealth of research supports this idea that contextual stability facilitates temporal integration of elements into coherent memories. Further, empirical studies demonstrate that a wide variety of context changes, such as shifts in space^6^, emotion^7^, goals^8–13^, or perceptual features^14–16^, lead to the separation of adjacent memories. While these behavioral effects are robust and replicable, little is known about the mechanisms triggered at boundaries that adaptively segment and encode unique new memories.

The hippocampus plays a critical role in binding sequential or temporal associations essential to episodic memory^4,17–22^. Past research demonstrates, a constant push-and-pull between hippocampal encoding and retrieval mechanisms that promotes the storage of distinct memories. When encountering new information, hippocampal operations should reflect whether to encode a distinct memory trace (i.e., requiring ‘pattern separation’^23,24^) or incorporate these new details into existing memory representations (i.e., requiring ‘pattern completion’^25,26^). Importantly, hippocampal activation also appears to be sensitive to event structure. Across a wide range of paradigms, hippocampal activation and hippocampal-cortical connectivity at event boundaries relate to successful encoding and consolidation of recent events^4,13,27–32^. Focusing on these hippocampal mechanisms at event boundaries, however, has led researchers to overlook a fundamental question: are there other neural signals that can tip the balance between hippocampal separation and integration processes to adaptively segment and encode unique memories?

One possibility is that the locus coeruleus (LC), the primary supplier of norepinephrine (NE) to the brain, is the origin and substrate of this neural signal, serving to ‘reset’ hippocampal memory representations during context shifts. Through its widespread projections, the LC facilitates arousal, attention, and memory processes during salient occurrences^33–36^. LC neuronal inputs are particularly dense to the dentate gyrus (DG) subfield of the hippocampus, a region implicated in pattern separation^24^. This specialized anatomy provides a pathway through which noradrenergic activity could amplify memory separation at boundaries and help to disambiguate representations of temporally adjacent contexts^37–39^. Phasic LC responses signal contextual novelty and aid in encoding new memories^35,40–42^. When expectations about unfolding experiences are violated, phasic LC responses help signal prediction errors that rapidly update mental models^46–4.43^. The resulting global release of NE is thought to initiate a ‘network reset’, whereby functional brain networks become reorganized to prioritize processing new information^44,45^. In everyday life, event boundaries punctuate and signal critical moments of change in the world. Consequently, our ability to understand unfolding experiences may depend on a rapid and behaviorally relevant updating signal from the LC when something surprising, unexpected, or important occurs.

Studies examining pupil dilation, a putative index of LC activity^46–51^, provide initial support for the idea that noradrenergic system engagement at boundaries helps to structure memory. Our previous work demonstrates that event boundaries reliably elicit pupil dilation^16^. Moreover, distinct temporal components of pupil dilations relate to behavioral correlates of event segmentation, including subjective time dilation and reduced temporal order memory for information spanning those transitions^16^. These correlational findings are also corroborated by studies that manipulate arousal states more directly. For example, both highly arousing emotional sounds and prediction error-related arousal elicit event segmentation during sequence encoding, as indexed by changes in temporal memory^7,52^. Whether these connections between arousal and memory organization reflect changes in LC-NE activation remains unclear.

Here, we combined high-resolution functional magnetic resonance imaging (fMRI) and pupillometry to investigate whether engagement of arousal and noradrenergic mechanisms relate to event segmentation during the well-validated Ezzyat-Dubrow-Davachi (EDD) Paradigm^16,53^. Our findings support the idea that noradrenergic mechanisms contribute to the adaptive structuring of memory. First, we replicate established behavioral evidence that context shifts elicit behavioral memory separation and pupil-linked arousal^16^. We also find evidence for the critical missing link between LC activation and memory separation: boundary-induced LC activation selectively correlates with order memory impairments across boundaries and not within events. We furthermore find that boundaries result in a change in neural patterns between event-spanning items in the dentate gyrus (DG), with DG pattern differentiation being predicted by heighted LC activation. Together, these task-related findings align with the notion of an LC-mediated ‘reset’ signal that functionally configures hippocampal networks to represent contextually distinct events. We also find that higher signal intensity in the LC in neuromelanin MRI images^4–66^, an indirect neurochemical measure of repeated stress-related activation of the noradrenergic system^54^ and hyperarousal^54,55,56^, relates to diminished pupil responses at boundaries.

Low-frequency, task-unrelated fluctuations in LC activation also relate to diminished pupil dilation and phasic increases in LC activation at boundaries during the task. Elevated trait-like levels of arousal might therefore constrain the transient LC responses that signal important changes in the environment and, in turn, facilitate encoding of distinct events.

## Behavioral Results

### Event boundaries increase response times and induce event segmentation effects in long-term memory

Participants studied lists of neutral objects while listening to simple tones played in their left or right ear (**Figure 1A**)^16^. Temporal stability and change in the surrounding auditory context were used to create the perception of stable auditory ‘events’ and ‘event boundaries’, respectively. Eight pure tones were repeated in the same ear to create a sense of contextual stability. However, after 8 successive items, the tone switched to the other ear and changed in pitch to elicit perception of an auditory context shift, or event boundary. The new tone/ear then remained the same for the next 8 items before switching back again and so on.

**Figure 1.**
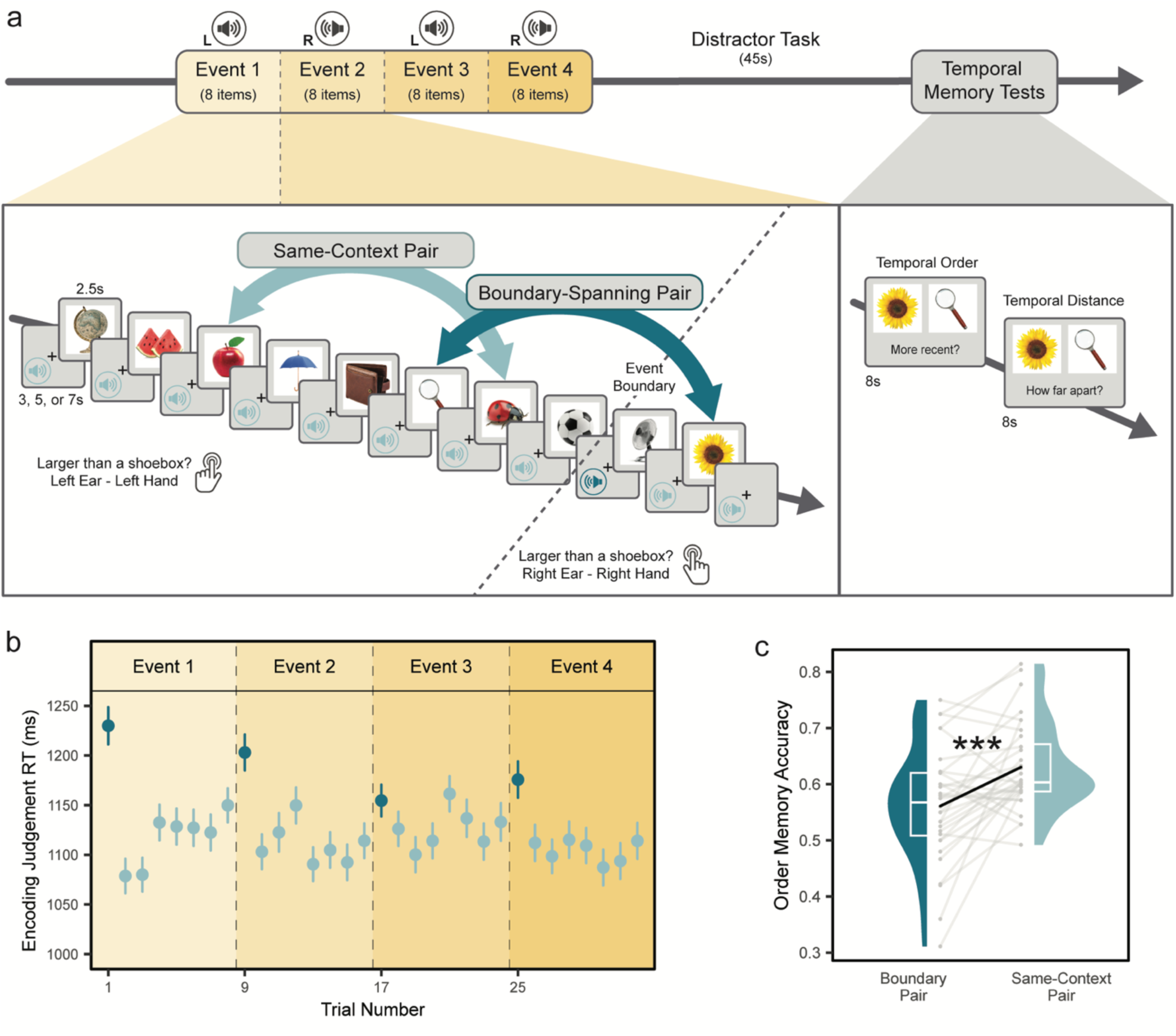
Auditory event boundaries elicit increased attention and the segmentation of distinct events in memory. (A) In the event sequence task, participants studied slideshows of 32 neutral object images.

We first compared response times to boundary items and same-context items for the object size judgements as an index of attention (**Figure 1A**). Participants were slower at judging objects that appeared immediately after a tone switch (*M* = 1178ms, *SD* = 276) compared to items that appeared after a repeated, same-context tone (*M* = 1115ms, *SD* = 286; t(7886) = 6.39, ß = 0.12, p < .001, 95% CI [0.08, 0.15]; **Figure 1B**). This response-time slowing effect most likely reflects a cognitive switching cost rather than a motor switching cost, as participants had ample time to switch hands during the tone cues (range: 1.5-3.5 seconds from tone to response; also see^10^ for similar argument).

Next, we tested if tone switches influenced later memory separation, as revealed by their effects on temporal order memory. To-be-tested item pairs were selected to have the same of number of intervening items and the amount of time was identical across all these pairs: each pair had three items between them and the time between them was always 32.5 seconds. Keeping this spacing and timing fixed allowed us to isolate the effects of context changes on memory. We also chose to include three intervening items, because this spacing allowed us to test the maximum number of pairs without repeating any items during the memory test (**Figure S1)**.

We specifically focused on temporal order memory effects, given predictions that noradrenergic system activation primarily elicits segmentation by disrupting temporal binding processes^57^. Replicating prior work, we found that temporal order memory was significantly impaired for boundary-spanning item pairs relative to same-context item pairs, a behavioral index of memory separation (ß = -0.15, p < .001, 95% CI [-0.21, -0.08], odds ratio = 0.86; **Figure 1C**).

Interestingly, we also found that increased attention at boundaries, indexed by slower RTs, predicted larger impairments in temporal order memory across participants. This trade-off effect suggests that greater local processing at event transitions occurred at the expense of maintaining temporal encoding processes across time^15^. We report these results below (see “**Individual Differences Correlations between Pupil-Linked Arousal, LC Measures, and Behavior”).** In summary, our behavioral results accord with the idea that boundaries trigger attention and support the temporal organization of events in memory.

Participants heard a pure tone in either their left or right ear before each image, which signaled which hand they should use to judge the size of each object. After 8 successive items, the tone switched to the other ear and repeated for the next 8 items, switched back, and so on. Thus, repeated tones created a stable auditory event, whereas tone switches created ‘event boundaries’ that divided the sequences into four events.

Following each list, participants performed two temporal memory tests used to operationalize event segmentation: temporal order memory and temporal distance ratings. (B) Responses times (RTs) for the object judgements plotted by item position in the lists. Dark blue colors indicate boundary items, or objects following a tone switch. Light blue items indicate objects that followed a repeated tone. Dots represent mean RTs. Vertical solid bars represent standard errors of the mean (s.e.m.). (C) Event boundaries impaired later temporal order memory for item pairs that spanned a boundary compared to pairs encountered within the same auditory context. Colored boxplots represent 25th–75th percentiles of the data, the center line the median, and the error bars the s.e.m. Individual dots represent individual participants (*n* = 32). Statistical results reflect results of a logistic mixed effects model. ***p < .001.

## Effects of Event Boundaries on Locus Coeruleus (LC) Activation

### LC activation at event boundaries relates to subsequent impairments in temporal order memory, a behavioral index of memory separation

Turning to our key hypotheses, we examine if LC activation at event boundaries (i.e., tone switches) relates to later memory separation. LC activation was quantified using “parameter estimates”, or beta values representing the activation levels of LC voxels to each tone during encoding. To simplify, we use the term “LC activation” to describe stimulus-evoked LC responses. These estimates of LC activation were extracted using hand-drawn anatomical masks derived from each participant’s LC MRI images (i.e., neuromelanin-sensitive scans; see **Figure S2** for study-specific mask).

Overall, there was no significant modulation of LC activation either by boundary tones (*M* = -1.75, *SD* = 717.59; t(848) = -0.07, p = .94; 95% CI [-50.03, 46.54]) or by same-context tones (*M* = 3.12, *SD* = 663.26; t(8222) = 0.36, p = .72; 95% CI [-13.05, 18.96]) relative to baseline (**Figure 2A**). We also did not find a significant difference in LC activation between conditions (t(9072) = -0.20; ß = -3.63e-03, p = .84, 95% CI [-0.04, 0.03]), suggesting that, on average, boundaries do not always elicit a strong LC response. Trial-level responses across the sequences are displayed in **Figure S3.**

**Figure 2.**
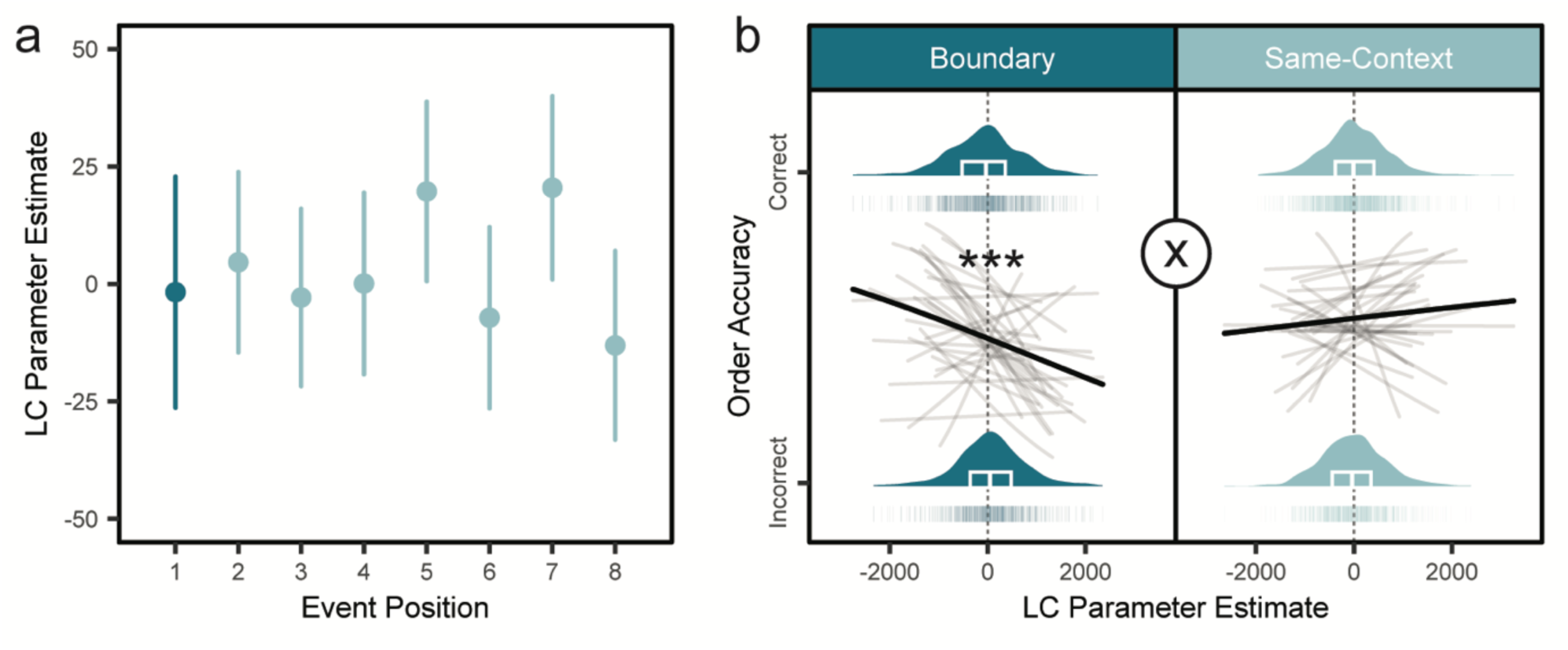
Locus coeruleus (LC) activation at event boundaries relates to impaired temporal order memory. (A) Mean LC parameter estimates, a measure of BOLD signal or level of brain activation, to each tone during encoding plotted as a function of item position within the 8-item auditory events. Boundary-related LC responses (item position #1 within an event) are displayed in dark blue, whereas repeated tone-evoked LC responses (item positions #2-8 within an event) are displayed in light blue. Vertical bars represent s.e.m. (B) Trial-level logistic regression between tone-evoked LC activation and temporal order memory accuracy, separated by condition. Dark, bold lines represent the average regression slope across all participants. Light grey lines represent regression slopes for each participant from the logistic regression. Plots display density distributions and colored boxplots represent 25th–75th percentiles of the data, the center line the median, and the error bars the s.e.m. The data includes *n* = 32 participants. ***p < .001 main effect for boundary trials. The “X” symbol indicates significant LC-by-condition interaction effect on order accuracy with p = .002.

Although boundaries did not reliably activate the LC across the task, we were most interested in seeing whether trial-level engagement of the LC was related to the temporal order memory impairments observed at boundaries. Consistent with this core prediction, logistic mixed effects modeling revealed a significant condition-related interaction effect of LC activation on temporal order memory, such that tone-evoked LC responses were significantly more coupled with order memory impairments on boundary trials compared to same-context trials (ß = -0.11, p = .002; 95% CI [-0.18, -0.04], odds ratio = 0.90, **Figure 2B**).

When examining the two conditions separately, we found that this interaction effect was driven by trial-level boundary-induced LC activation relating to larger impairments in temporal order memory (ß = -0.18, p < .001; 95% CI [-0.28, -0.07]; odds ratio = 0.84; **Figure 2B**). This LC-memory relationship was not seen for same-context item pairs (ß = 0.04, p = .39; 95% CI [-0.05, 0.13]; odds ratio = 1.04, **Figure 2B**). Thus, these results show that activation of the noradrenergic system at boundaries relates to disruptions in the sequential integration of items in long-term memory.

## Hippocampal Pattern Similarity fMRI Analyses

### Event boundaries promote the temporal differentiation of activation patterns in left hippocampal dentate gyrus, while also increasing pattern similarity in left CA2/3

Thus far, we have shown that LC activation at boundaries relates to later memory separation. Next, we examined whether boundaries decrease the temporal stability of multivoxel activation patterns, an index of pattern separation, in a region sensitive to noradrenergic activity: the hippocampal dentate gyrus (DG).

First, we extracted encoding-related multivoxel activation patterns from left and right hippocampal subfields (CA1, CA2/3, and DG) for the two images that would be subsequently queried for temporal memory (see **Figure S4** for an example of one participant’s segmentation). These multivoxel patterns were then correlated between each of the to-be-tested item pairmates, providing a trial-level measure of hippocampal subfield pattern similarity. Because different hippocampal sub-regions make both overlapping and unique contributions to episodic memory processes^24,25^, we modeled all six of the subfields’ pattern similarity estimates as fixed effects predictors of Condition in a multiple logistic regression, which was modeled as a binary outcome variable (boundary = 1, same-context = 0). For all multiple regression analyses, the VIF’s were less than 1.5, verifying low collinearity between predictors.

We found that item pairs spanning an event boundary were associated with lower pattern similarity, or increased pattern separation, in left DG (ß = .09; p = .035; 95% CI [6.33e-03, 0.18]; odds ratio = 2.58; **Figure 3, bottom left**). Boundaries also led to a marginally significant increase in right CA2/3 pattern similarity (ß = -0.08, p = .057; 95% CI [-0.16, 2.57e-03], odds ratio = 0.51). There were no other statistically significant effects in the other four subfields (all p’s > .10; **Figure 3, bottom left).**

**Figure 3.**
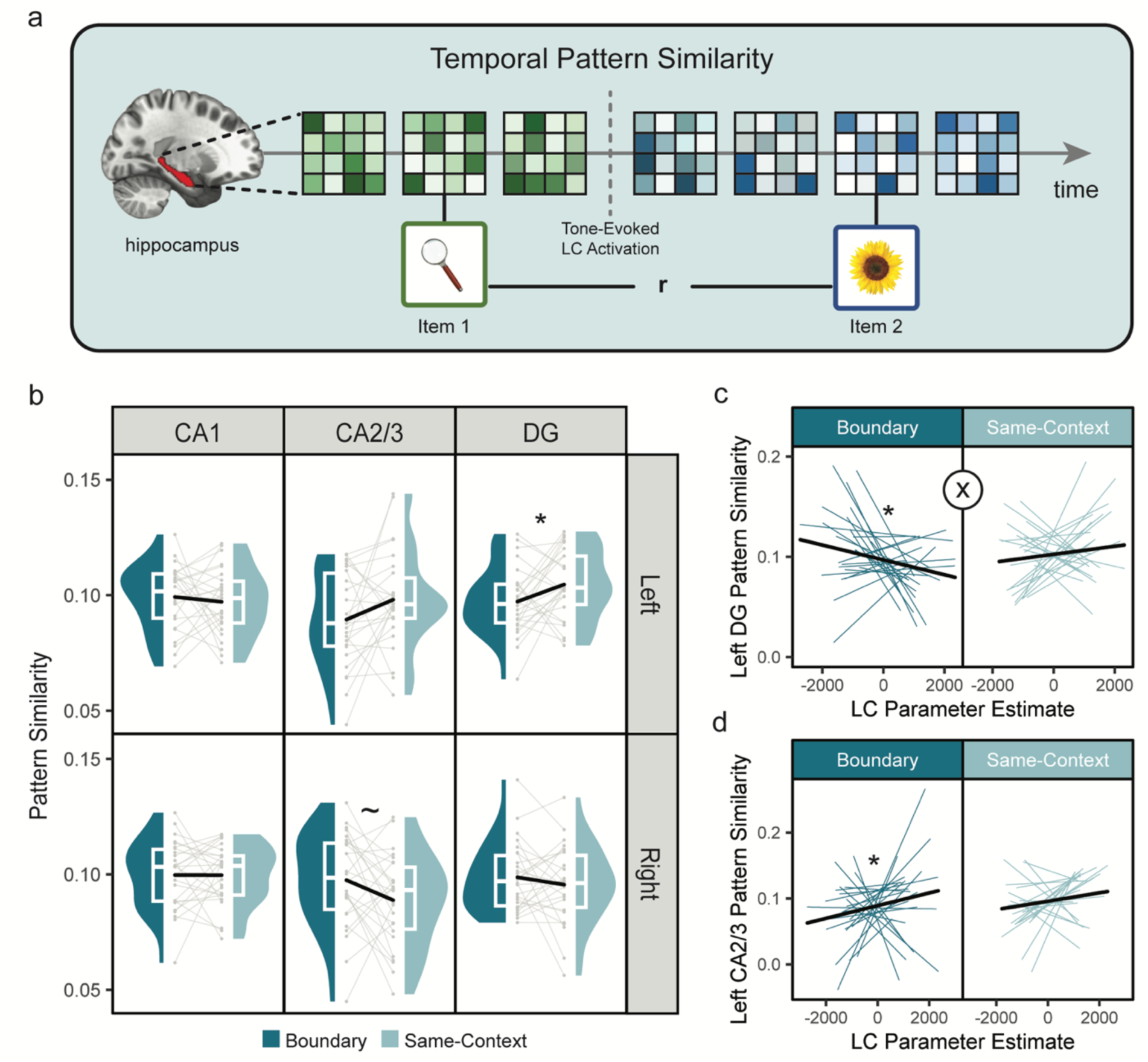
Event boundaries led to more differentiation of left dentate gyrus (DG) multivoxel activation patterns across time, and this pattern separation effect corresponded with heightened LC activation at those same boundaries. (A) Schematic of key hypothesis that LC activation at event boundaries predicts greater temporal pattern separation in hippocampal DG activation patterns. Colored grids represent multivoxel activation patterns that were extracted from each hippocampal subfield for the to-be-tested item pairs. Subfield pattern similarity was quantified as the linear correlation (r) between these vectorized multivoxel patterns of activation during encoding. (B) Boxplots showing differences between hippocampal subfield similarity for boundary pairs versus same-context item pairs during encoding. Colored boxplots represent 25th–75th percentiles of the data, the center line the median, and the error bars the s.e.m. Overlaid dots and connecting grey lines represent data from individual participants (*n* = 25). Note that statistical significance reflects the outcome of multiple logistic mixed effects modeling analyses, where all six subfields’ pattern similarity values were mean-centered by participant and simultaneously modeled as fixed-effect predictors of Condition. (C) Linear mixed effects modeling results demonstrating that greater trial-level LC activation at event boundaries predicts the degree of pattern dissimilarity in left DG specifically, a neural measure of pattern separation. By contrast, intervening LC activation between to-be-tested same-context pairs did not significantly predict left DG pattern similarity for those same item pairs. (D) The results also revealed a significant main effect of LC activation on pattern similarity in left CA2/3, with greater pattern similarity driven by boundary trials. Individual colored lines represent regression slopes for each participant (*n* = 25). Dark, bold lines represent the average regression slope across all participants. The “X” symbol represents a statistically significant LC-by-condition interaction effect with p = .02. *p < .05; ∼p < .10.

In a similar style of multiple regression analysis, we examined if hippocampal multivoxel patterns were related to temporal order memory for those same item pairs, and whether those correlations differed by Condition.

There were no statistically significant main effect or condition-related interaction effects between any hippocampal subfield and temporal order memory, suggesting that, within the hippocampus, boundaries primarily promoted neural and context differentiation rather than memory changes (all p’s > .10).

### Increased LC activation at event boundaries relates to opposite effects on pattern similarity in left DG and left CA2/3

In the next set of linear mixed effects modeling fMRI analyses, we tested another key hypothesis that boundary-induced LC activation promotes neural differentiation in left DG representations, given the critical role of LC-DG pathways in encoding distinct episodic memories (**Figure 3, top panel**). As before, we modeled all six of the subfields’ pattern similarity estimates as fixed-effect predictors of tone-related LC activation, in addition to a main and interaction effect of Condition.

For boundary trials, we found that greater LC activation at boundaries was related to more dissimilar left DG patterns between item pairs that spanned those same boundaries (t(1086) = -2.41; ß = -0.08, p = .016, 95% CI [-0.15, -0.02]; **Figure 3C**). By contrast, there was not a significant LC-DG correlation for same-context pairs (t(1447) = 0.66; ß = 0.02, p = .51; 95% CI [-0.04,0.08]; **Figure 3C**). Importantly, we also found a significant condition-by-LC activation interaction effect, such that LC activation was more correlated with lower pattern similarity in left DG for boundary-spanning pairs compared to same-context pairs (t(2535) = -2.33; ß = -0.05, p = .020, 95% CI [-0.10, -8.22e-03]; **Figure 3C**).

LC activation also elicited a significant increase in left CA2/3 pattern similarity across conditions, (t(2535) = 2.46,; ß = 0.05, p = .014; 95% CI [0.01, 0.09]). This LC-CA2/3 association primarily emerged on boundary-spanning trials (t(1086) = 2.15; ß = 0.07, p = .032; 95% CI [5.94e-03, 0.13]; **Figure 3D)** and was not observed on same-context trials t(1447) = 0.91; ß = 0.03, p = .36; 95% CI [-0.03,0.08]). However, there no condition-related interaction effects in LC-CA2/3 coupling, so we cannot draw strong interpretations about the selectivity of these effects at boundaries (t(2535) = 1.17; ß = 0.02, p = .24; 95%; CI [-0.02, 0.07]; **Figure 3D**). Finally, there were no other significant main, condition-specific, or condition-related interaction effects of tone-evoked LC activation on pattern similarity in the remaining subfields (all p’s > .05).

Together, these results demonstrate that boundary-evoked LC responses relate to greater temporal pattern separation in left DG. Moreover, this functional reconfiguration effect of LC activity was specific to boundaries and the left DG subfield, suggesting that separation mechanisms are only engaged during behaviorally relevant moments when differentiation is needed. At the same time, LC activation was related to the increased stabilization of left CA2/3 patterns, which was driven by boundary trials but not exclusively.

## Pupillometry Results

### Distinct pupil dynamics are sensitive to event boundaries

Prior work shows that boundaries trigger pupil dilation and engage central arousal processes^16^. Pupil dilation, however, is complex and mediated by multiple autonomic pathways and neuromodulatory systems^47,58^. Building on earlier work, we use a temporal principal component analysis (PCA) to decompose boundary-related pupil dilations into its distinct temporal features, providing a way to link event segmentation to noradrenergic processes. We also use this PCA approach to examine whether more general pupil-linked arousal mechanisms relate to memory organization and attention.

We first compared average mean pupil dilation responses during tone switches (boundary tones) to mean pupil dilation responses to the repeated tones (i.e., tones preceding items 2-8 within any given 8-item auditory event (**Figure 4A and Figure 4B**). As expected, boundaries elicited significantly larger pupil dilations (*M* = 118.95, *SD* = 201.46) than same-context tones (*M* = 3.21, *SD* = 182.13; t(7347) = 15.96; ß = 0.31, p < .001; 95% CI [0.27, 0.35]; **Figure 4A and 4B**).

**Figure 4.**
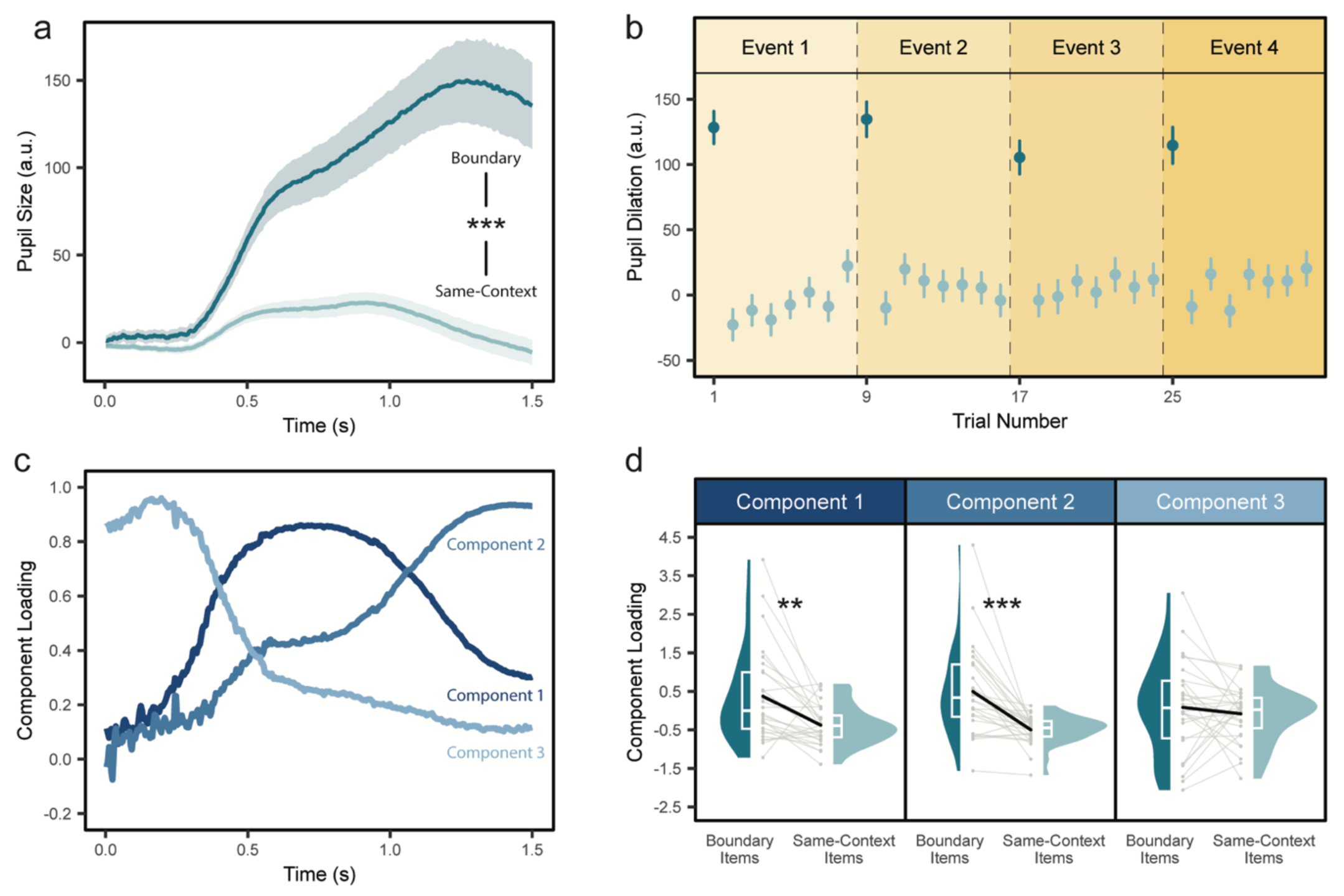
Event boundaries influence distinct temporal characteristics of pupil dilation. (A) Time-course showing the average pupil dilation response evoked by boundary tones (dark green), or tone switches, and same-context, or repeated, tones (light blue). Shaded windows represent s.e.m. at each timepoint. (B) Average pupil dilations plotted as a function of item position within the sequences. Dark blue colors indicate boundary items, or those objects that immediately followed a tone switch. Light blue items indicate objects that immediately followed a same-context, or repeated tone, within a given 8-item auditory event. Vertical bars represent s.e.m. (C) Three temporal features of tone-evoked pupil dilations identified using a temporal principal components analysis (PCA). (D) Statistical comparisons between PCA loadings of the three pupil components between boundary and same-context items. Colored boxplots represent 25th–75th percentiles of the data, the center line the median, and the error bars the s.e.m. Grey dots and connecting lines indicate datapoints from individual participants (*n* = 28). **p < .01; ***p < .001.

Next, a temporal PCA was used to decompose average pupil dilation into its constituent temporal features^16^. All pupil samples were averaged across the time-window of tone-evoked pupil dilations (i.e., onset of tone plus 1.5 seconds) and across participants. The PCA revealed three canonical pupil components identified in prior work, including a biphasic response that may index separate influences of parasympathetic and sympathetic nervous system regulation on pupil diameter^16,59^ (**Figure 4C**). The temporal characteristics of these pupil components, including their latencies-to-peak and percent of explained variance, were as follows: (1) an early-peaking component (684 ms; 89.26% variance); (2) intermediate-peaking component (1,420 ms; 8.40% variance); and (3) a slowly decreasing component (19.6 ms; 1.27% variance).

Using paired t-tests, we found that boundaries significantly modulated loadings, or engagement, of the early-peaking pupil component (component #1; t(27) = 3.63, p = .001, Cohen’s d = 0.69; **Figure 4D**) and the intermediate-peaking pupil component (component #2; t(27) = 4.76, p < .001, Cohen’s d = 0.90; **Figure 4D**). In contrast, there was no significant effect of event boundaries on loadings for the slowly decreasing pupil component (component #3; t(27) = 0.67, p = .51, Cohen’s d = 0.13; **Figure 4D**).

## Individual Differences Correlations between Pupil-Linked Arousal, LC Measures, and Behavior

Using an individual differences approach, we next examined relationships between LC structure/function, attention, and memory separation. For all measures (except for LC MRI contrast-to-noise ratio), we computed a difference score by subtracting average values for same-context trials from the average values for boundary trials to isolate the specific effects of context shifts. We then correlated these variables across participants using Spearman’s rank order correlation analyses.

### An early-peaking component of pupil-linked arousal relates to temporal order memory impairments and response-time slowing at boundaries

First, we investigated whether pupil-linked arousal was related to behavioral metrics of event segmentation. We found that individuals who showed greater boundary-driven engagement of the early-peaking pupil component also exhibited greater impairments in temporal order memory across boundaries (component #1; ρ = -0.45, p = .016; **Figure 5A**). No other pupil-memory associations were observed for the other two pupil components (all p’s > .36).

**Figure 5.**
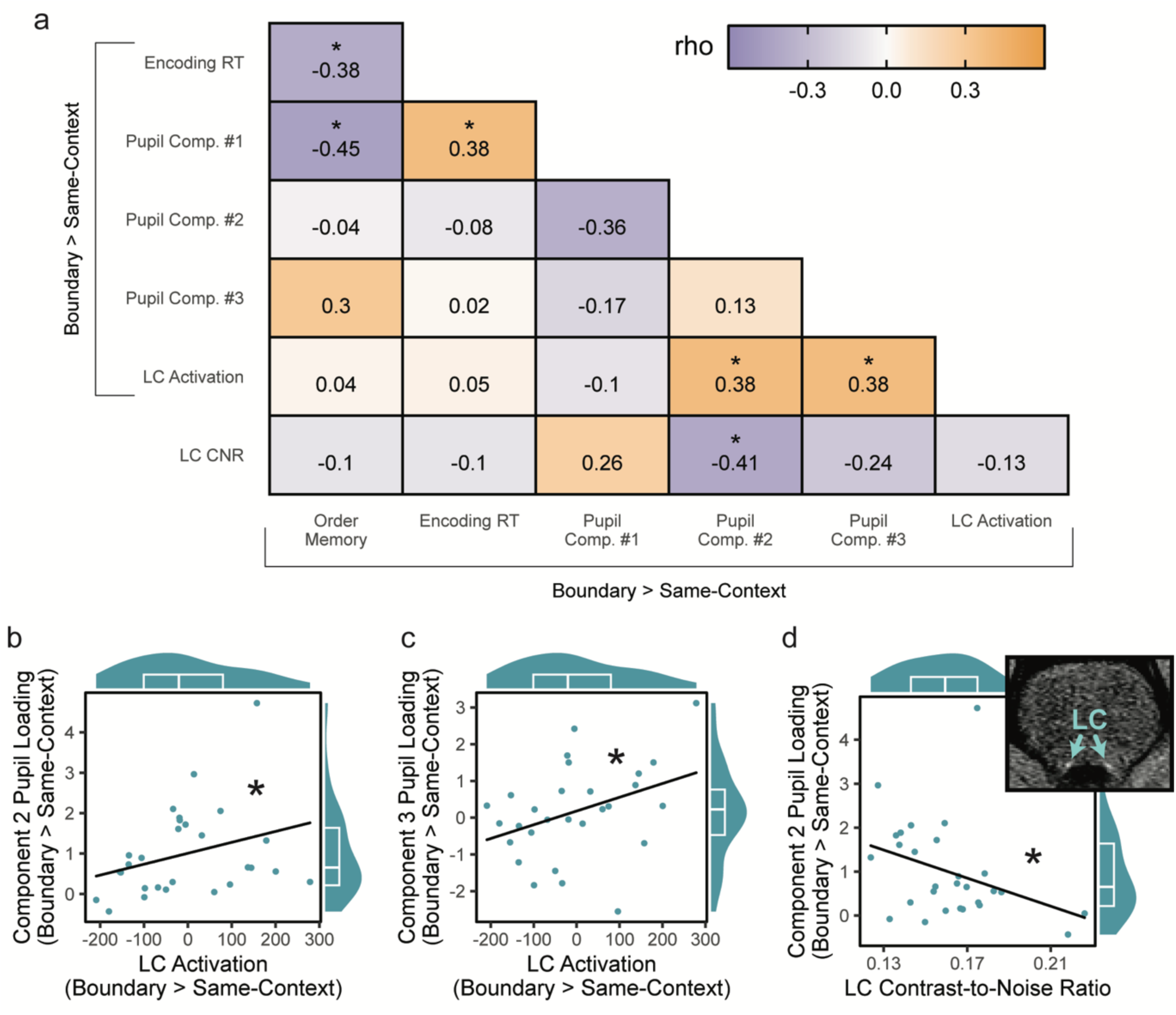
Pupil-linked arousal relates to patterns of LC activation at event boundaries and a neurochemical index of LC structure. (A) A Spearman’s rank order cross-correlation matrix relating key neurophysiological measures of arousal, behavioral metrics of memory separation, and an MRI measure of LC structure across participants. For all functional and behavioral measures, we used subtraction scores of boundary trials versus same-context trials to isolate boundary-specific effects. Pupil variables included loadings from the three aspects of pupil dilation identified by the temporal PCA. Noradrenergic activity was assessed using tone-evoked changes in LC activation. Behavioral metrics included response-time (RT) slowing at boundaries and boundary-related temporal order memory impairments. LC structure was assessed by extracting the LC contrast-to-noise ratio (CNR) in the fast spin echo images. Within the correlation matrix, colored bar indicates the Spearman’s rho correlation coefficient, with purple values indexing negative correlation coefficients and orange boxes indexing positive correlation coefficients. Color saturation reflects the strength of the correlations. (B/C) Spearman’s rho correlation plots showing that higher boundary-induced LC activation was positively correlated with greater boundary-related engagement of pupil dilation component #2 and #3, respectively. (D) Spearman’s rho correlation plot showing that higher LC CNR was associated with smaller boundary-related engagement of pupil dilation component #2. For all correlations (panels B-D), marginal density-boxplots depict the median (center line) and interquartile range (25th–75th percentiles) of the x- and y-distributions, and individual dots represent each participants’ data. (D) An example participant’s fast spin echo MRI image, a structural sequence sensitive to neuromelanin signal or cellular water content. Bilateral LC nuclei appear as bright dots that standout from neighboring brainstem tissue and ventricles (arrows). All behavioral and brain correlations have *n* = 32 participants; pupil-related correlations have *n* = 28 participants. *p < .05.

Boundary-driven engagement of this early-peaking pupil component was also positively correlated with greater response time (RT) slowing at boundaries, potentially indexing enhanced attention during context shifts (component #1; ρ = 0.38, p = .016; **see Figure 4C for component**). These results suggest that a pupil dilation component previously implicated in task-relevant responses and parasympathetic nervous system activation^59,60^ relates to disruptions in temporal binding across boundaries as well as a boost in attention. We furthermore found that boundary-related RT slowing was correlated with larger boundary-related impairments in temporal order memory (ρ = -0.38, p = .031; **Figure 5A**). Prioritizing local information at event transitions therefore appears to disrupt sequential integration processes^15^.

### Two putative sympathetic nervous system components of pupil-linked arousal relate to LC activation at boundaries

We next aimed to link a specific sympathetic nervous system-related pupil component, which peaks ∼1.5s post-stimulus^60^, to phasic increases in LC activation at boundaries (**see Figure 4C for component)**. Our analyses indeed revealed a significant positive correlation between LC activation evoked by boundaries and loadings on both the intermediate-peaking pupil component (component #2; ρ = 0.38, p = .044; **Figure 5A and 5B**) and slowly decreasing component (component #3; ρ = 0.38, p = .043; **Figure 5A and 5C)**. In contrast, boundary-related LC activation was not significantly correlated with the remaining pupil component that peaked around 700ms (p > .05). Thus, two distinct temporal features of stimulus-evoked pupil dilation appears to specifically capture noradrenergic effects, as the LC regulates sympathetic outflow^61^.

### LC contrast-noise (CNR), a potential indirect measure of background or sustained LC activation, relates to diminished boundary-evoked pupil dilations across participants

In a final across-participant correlation analysis, we tested if a trait-like metric of sustained LC output, LC CNR from fast spin echo MRI images, relates to changes in pupil dilation and LC activation at boundaries (**Figure 5D**). Spearman’s rank order correlations revealed that higher LC CNR was anti-correlated with boundary-related loadings on the intermediate-peaking pupil component (component #2; ρ = -0.41, p = .033, **Figure 5D** and **5C; see Figure 4C for component**), consistent with the idea that tonic LC activity blocks task-induced phasic LC activity^62^. LC CNR was not significantly correlated with boundary-related loadings on the other two pupil components (both p’s > .05). In contrast to the pupil findings, higher LC CNR was not significantly correlated with boundary-induced LC activation itself (ρ = -0.13, p = .47; **Figure 5A**).

### Structural and functional estimates of elevated background LC activation relate to reduced boundary-induced pupil dilations and LC responses across participants

To bolster our interpretation that LC CNR captures chronic patterns of LC activation, we performed a series of exploratory analyses targeting low frequency BOLD signal fluctuations as a proxy for tonic LC activity (see **Figure S5A**). We reasoned that slower, task-unrelated fluctuations in LC activation may index more sustained, background changes in arousal and attention, as suggested by prior resting-state fMRI work^63^.

Our results support the notion that higher tonic LC activation can mask transient, task-relevant responses (see **Figures S5B-F)**. Across participants, higher LC low-frequency (LF) BOLD variability (0.1-.01Hz range) was positively correlated with LC CNR (ρ = 0.32, p = .037, one-tailed). Higher LC-LF BOLD variability was also anti-correlated with boundary-related engagement of pupil component #2 during the task (ρ = -0.34, p = .04, one-tailed). Additionally, higher power in this frequency domain was significantly anti-correlated with event boundary-related engagement of pupil component #2 (ρ = -0.33, p = .046, one-tailed) as well as smaller boundary-related LC activation during the task (ρ = -0.54, p < .001, one-tailed). Together, these findings demonstrate that both structural and functional measures of hyperarousal and elevated background LC activation related to impaired event perception or salience detection.

## Discussion

The locus coeruleus (LC) is a core hub of the arousal system that facilitates attention and memory^35^. Discrete, momentary bursts of LC activity are essential for signaling novelty and shifts in environmental contingencies, which could be construed of as ‘event boundaries’ that punctuate continuous experience. Here, we provide strong evidence that pupil-linked arousal and LC signaling facilitate organizational changes in episodic memory. We first replicated findings that boundaries segment contextually distinct memories and elicit pupil-linked arousal responses. Importantly, we now find evidence for a critical mechanistic link between LC activation at boundaries and temporal pattern separation in left dentate gyrus (DG). We propose that this neural separation effect could signify a mental reset of ongoing mental representations, mediated by an LC signal during event transitions. Additionally, indirect functional and structural measures of hyperarousal were associated with impaired event processing, suggesting that event segmentation is shaped by complex interactions between tonic and phasic patterns of noradrenergic activity.

### Locus coeruleus engagement at event boundaries selectively relates to later memory separation

Our key novel finding was that increased LC activation at boundaries was significantly correlated with reduced temporal order memory across those events, a common behavioral marker of memory separation. This important finding aligns with many influential theories of LC function previously only tested in animal models, including its proposed role in resetting functional brain networks during shifts in environmental contingencies or surprising moments^44,45^. For over a decade, researchers have speculated that catecholamines drive event segmentation processes, largely due to their ability to broadcast prediction errors across the brain and to coordinate attentional and memory processes^64–66^. Using converging methods of pupillometry, fMRI, and behavioral measures of memory separation, we demonstrate that LC engagement during salient context shifts predicts event segmentation effects. Importantly, this LC-memory relationship was specific to boundary trials and was not observed within stable auditory events. LC engagement thereby only influences memory separation during behaviorally relevant moments when internal context representations are rapidly updated and encoded as novel events. If such event model-updating processes were triggered by *any* phasic LC response, memories would potentially become fragmented and organized inappropriately, as they would not align with meaningful environmental changes.

### Pupil-linked arousal and locus coeruleus activation relate to the memory-parsing effects selectively at event boundaries both within and across individuals

In the present study, event boundaries did not significantly modulate the average LC response. This suggests that not all context transitions are equally meaningful and warrant memory separation. One possibility for this null effect is that our boundaries simply were not salient enough to always enlist the LC. Event segmentation is differentially engaged based on task demands and the goal relevance of event boundaries^4,8,10^. As such, simply presenting salient arbitrary stimuli, even if novel, does not necessarily constitute a boundary in long-term memory. For example, simple target detection, which elicits pupil dilation^67^, does not relate to impaired order memory, which is used to operationalize event segmentation^68^. The LC might thereby only drive memory separation when there are meaningful changes in the structure and statistics of the environment that violate an active, stable model of ongoing events^57,69^.

Our individual difference results furthermore suggest that some individuals may simply be more sensitive to the presence of event boundaries. We found positive associations between boundary-related LC activation and two specific temporal features of pupil dilation. One of these pupil components exhibited a late peak, typically associated with mental resource allocation^60^ and sympathetic nervous system activation^60,61,70^. The other pupil component, which peaked much earlier, is thought to reflect pre-stimulus anticipatory processes^16,71^. This pre-stimulus pupil effect suggests that some individuals may recruit LC-related processes to predict and segment memories proactively. Indeed, stronger loading on this pupil component has been shown to be related to greater memory separation in a version of this experiment where stimulus timings were fixed^16^. This pupil-memory effect disappeared in the current study, where the only difference was that stimulus timings were jittered and thus unpredictable. A more proactive segmentation strategy could therefore be less effective when the precise timings of event transitions are unknown. Future studies could test this hypothesis more directly by varying the predictability of event structure within the same task.

The PCA analysis also revealed a separate pupil response that peaked around 700 milliseconds that was correlated with memory separation and a slowing of event boundary response times. These findings suggest that this intermediate-peaking pupil component may be capturing a known trade-off that occurs between enhanced local boundary processing and ongoing temporal encoding processes^15^. Prior work has linked this pupil component to motor and decision-related processes as well as parasympathetic nervous system regulation^60,61^. One possibility is that it captures contributions from cholinergic activity, as this brain system supports parasympathetic control over pupil dilation^47^ as well as enhanced attention and item encoding^72^. Another possibility is that response-time slowing at boundaries is relate to the cognitive cost of task switching^10^. Alternatively, or in combination, this slowing effect may index the increased processing demands of updating an event model^73,74^. In support of this idea, loading on this intermediate-peaking pupil component has been linked to motor remapping at event boundaries^16^, which captures the task-relevant audio-motor features that define distinct events in this task.

### Locus coeruleus activation may facilitate processing of novel events via inputs to left dentate gyrus and left CA2/3

Paralleling the effects of LC on behavioral measures of memory, we found that boundaries perturb ongoing contextual representations in the hippocampus, in part through activating the LC. Interestingly, this finding aligns with recent evidence in rodents showing that there is elevated release of norepinephrine in the hippocampus following putative event boundaries, an effect that appears to relate to more distinctive hippocampal representations of space^75^. Using multivoxel pattern similarity analyses, we showed that neural activation patterns in left DG were more differentiated for items with an intervening boundary compared to equidistantly spaced items pairs encountered within the same context. The extent of boundary-related DG temporal pattern separation corresponded with greater LC activation at those intervening boundaries. Like the LC-memory association, this LC-DG effect only occurred for boundary trials and not when the LC was arbitrarily engaged within a stable event.

This functional specificity of LC-DG modulation is highly adaptive because it provides a mechanism by which neural patterns shift during behaviorally relevant moments like boundaries. Consistent with prior frameworks of LC function, LC activation during salient context shifts may directly modulate DG neural patterns to reset ongoing temporal integration processes in hippocampal circuits, leading to the separation of sequential events^39^. Indeed, the LC is densely connected to the DG^37,76,77^ and regulates synaptic plasticity in this region^37,38,78^. In rodents, phasic LC activation has been shown to promote global remapping effects in DG, such that novel spatial maps are observed in familiar spatial environments^38,39^. The current work builds on these findings by showing that phasic LC activity also promotes the functional reconfiguration of DG representations for different temporal and perceptual contexts. Interestingly, rodent work has also shown that LC activation promotes the stabilization of place fields in CA3 during single-trial learning of a novel context^79^. We found that LC activation is related to higher left CA2/3 pattern similarity, perhaps a parallel to this earlier work. It is noteworthy that this LC-CA2/3 effect was mostly strongly coupled on boundary trials. However, because there was not a significant condition-related interaction effect, we cannot conclude that LC activation specifically modulates CA2/3 pattern stability during shifts in context. One possibility is that this pattern stabilization process persists into a new event, capturing higher-order representations of event structure in the sequences^80^. Future work in humans should aim to disentangle these distinct contributions of LC activation to context representations in CA2/3 and DG.

While we did not have an *a priori* hypothesis about laterality, we also found the boundary-related DG pattern separation effect was specific to the left hemisphere. Evidence of strong lateralization effects in hippocampus with respect to memory function is relatively sparse. Interestingly, however, recent work in rodents has shown that context discrimination is higher between spatial environments in the left versus the right DG^81^. Likewise, neuroimaging work in humans has recently linked dynamic patterns of left DG activation to event structure^80^. Future research should investigate the lateralized influences of different hippocampal processes to representing stability and change in the environment.

Our findings contribute to a growing body of research in humans investigating whether the LC can also facilitate a ‘network reset’ in distributed brain networks, driving sudden shifts in conscious awareness, effortful behavior, or salience processing^82–85^. For example, one study combined pupillometry, fMRI, and graph theoretical analyses to show that cognitive load-dependent increases in pupil diameter correspond to change in the activity and topography of large-scale brain networks^85^. Similarly, neuroimaging markers of phasic LC activation have been linked to task-related perceptual switches, which coincide with shifts in the brain’s “energy landscape”^84^. However, like foundational models of event segmentation, most neuroimaging studies have focused on an LC-driven functional reconfiguration of large-scale brain networks under arousal. Our results suggest that such attentional reorienting at event boundaries may also influence the functional organization of hippocampal networks, enabling the hippocampus to update its representations in response to changes in context. The theoretical LC-mediated network reset might therefore be a more general principle of adaptive cognitive processing, because it can coordinate task-relevant updates across many attentional- and memory-related brain networks.

### Indirect structural and functional measures of elevated background LC activation relate to smaller boundary-related arousal responses, holding important implications for understanding and treating disorders of hyperarousal and memory

Using a combination of pupillometry and a specialized structural MRI sequence, we found a specific pupil signature of boundaries that correlated with an index of LC structure. Specifically, across individuals, engagement of a sympathetic-related component of pupil dilation was anti-correlated with LC MRI contrast, or signal intensity, a putative marker of chronic LC activation. This relationship may reflect the known trade-offs between tonic and phasic modes of LC activation^62^. Elevated tonic patterns of LC activation, as indirectly evidenced by higher LC signal intensity and patterns of task-unrelated low-frequency LC activation, may constrain the sensitivity of arousal systems to transient environmental changes, potentially impairing event model updating when it matters. Such impairments may be especially pronounced under stress. Work in rodents shows that restraint-induced stress can augment infraslow fluctuations in LC activity (0-0.5Hz), which covers the same frequency range that we analyzed in the current study^86^.

Interestingly, our exploratory fMRI analyses also revealed a novel association between higher LC signal intensity and low-frequency fluctuations in LC BOLD signal. Presently, however, the nature of this structure-function relationship – as well as the link between LC neuromelanin and tonic LC activity more broadly - is still unclear. It is thought that higher LC signal intensity could reflect elevated NE production following prolonged periods of hyperarousal, because the LC help regulate the stress response^87^. Indirect support for this idea comes from studies showing that slow-paced breathing interventions known to quiet sympathetic outflow lead to reductions in LC signal intensity in healthy young adults^56^. Further, it has been shown that LC signal intensity is higher in combat-exposed veterans with PTSD compared to those without PTSD^54^ and is correlated with reduced parasympathetic control over the heart^55^. Through this lens, our observation of an association between boundary-induced arousal and a potential noradrenergic marker of stress may have important implications for developing interventions in disorders where disturbances in arousal and memory function intersect. Indeed, deficits in event perception are ubiquitous across disorders marked by aberrant arousal and memory function, including attention deficit hyperactivity disorder^88^, Parkinson’s disease^89^, post-traumatic stress disorder (PTSD)^90–92^, and age-related dementia^93^.

Insofar as discrete LC responses are needed to signal event boundaries, excessively high levels of arousal and tonic LC activation would likely disrupt normal event segmentation processes. When LC tonic is high, task-relevant phasic LC responses are reduced due to trade-offs between these two modes of activity^62^. But these maladaptive trade-offs could be mitigated, and several strategies could be used to restore “healthier” modes of LC phasic activity to facilitate event cognition and memory. For example, slow-paced breathing^56^, vagus nerve stimulation^94^, pharmacology^95^, and simple exercise^96^ have all been shown to enhance LC phasic activity and boost cognitive abilities. Such arousal-related interventions could also be used to augment event segmentation training, a behavioral technique of enhancing the salience of boundaries to improve event detection and enhance long-term memory^97^. Crucially, our findings also highlight a brain system that is likely more accessible for intervention than the hippocampus, which can only engaged indirectly by stimulating cortical regions on the outer surface of the brain^98^.

### Potential limitations and important considerations for future studies on event segmentation

There are several limitations of the current study that warrant consideration. While we report strong correlational relationships, we cannot make strong claims about causality. The LC is also notoriously difficult to study in humans due to its small size and its susceptibility to cardiac pulsation artifact in fMRI images^99^. Yet, a rapidly growing neuroimaging literature has provided convincing evidence that LC activation can be accurately measured and meaningfully linked to pupil dilation and/or behavior^50,51,96,100–105^. We used several strategies to mitigate potential issues with studying the LC in humans. First, we used a neuromelanin MRI scan to accurately localize and delineate anatomical masks of the LC in each participant, increasing the spatial specificity of BOLD measurements. Second, we included physiological nuisance regressors in our fMRI analyses for signal derived from ventricles, including the fourth ventricle neighboring the LC. These additional variables should have helped control for noise related to cardiac and brainstem pulsation artifacts. Noisy single-trial estimates were also filtered from analyses using a by-participant boxplot outlier detection method, helping reduce any spurious signals driven by physiological artifacts. Third, we did not apply spatial smoothing to the functional images, helping avoid smearing the BOLD signal beyond the true anatomical boundaries of the LC. While this approach sacrifices some of the signal-to-noise ratio, it also affords better spatial localization of LC-related signal. Fourth, we performed all fMRI analyses in participants’ native functional space, avoiding the potential spatial misalignments that can plague group-level LC analyses^106^. Fifth, we used a high-resolution imaging sequence with an in-plane spatial resolution of 1.5mm. This voxel size is sufficient to measure activation in the small LC, which is ∼1-3mm wide and ∼15mm long^107^. Finally, we corroborated our LC findings using pupillometry, lending additional evidence of a strong connection between phasic LC responses and task-related pupil dilations (for a review, see^46^).

Another potential limitation was the relatively coarse spatial resolution of our high-resolution anatomical images, which could influence the accuracy of hippocampal subfield segmentation. Some researchers have cautioned against the use of this algorithm for segmenting T1-weighted 1-mm^3^ isotropic resolution due to difficulties visualizing some internal structures in the hippocampus^108^. However, increasing evidence suggests that subfield segmentations are both replicable and reliable across a variety of factors, including different scanners, sampling time intervals, and sample sizes^109,110^. Prior work has also reported high intra-class coefficients for segmentations generated from both longitudinal and cross-sectional processing streams in individuals with PTSD^111^. Another neuroimaging study also internally corroborated this algorithm’s T1-image segmentation with high-field T2-weighted data within the same participants^112^. Despite these findings, it is important to acknowledge that the Freesurfer 6.0 segmentation approach is atlas-based and does not necessarily reflect anatomical ground truth^113^. Thus, our results should be interpreted with caution.

To verify the accuracy of our hippocampal segmentations as best as possible, we applied a rigorous, data-driven quality control procedure. We also were very conservative about data inclusion and removed entire participants who violated any of the outlier criteria^114,115^. Although it is common to combine the CA2/3 and DG subfields into a single anatomical mask, we chose to keep these ROIs separate due to substantial evidence that the DG is especially sensitive to fluctuations in noradrenergic activity^37^. We also found that LC activation at boundaries elicited completely opposite effects on left CA2/3 and DG pattern similarity, suggesting that these subfields were distinguishable and made unique contributions to processing novel contexts. Finally, spatial smoothing was not applied to our functional images to avoid blurring activation patterns across different hippocampal subfields. Future work could strengthen and validate our hippocampal and LC findings by using sophisticated imaging tools like cardiac-gated fMRI and high-resolution 7T fMRI.

An important strength of the current study was the use of a highly controlled and well-validated experimental paradigm to study segmentation effects in memory. This simplicity helped to limit complex interactions with other factors that impact attention and memory function, such as fluctuating task demands, competing contexts, and the semantic relevance of contexts and concurrent memoranda. Nevertheless, many interesting open questions remain about whether the LC is a shared mechanism of event segmentation across multiple contexts and types of event boundaries. For example, recent work suggests that shifts in emotional states can drive memory separation effects and influence temporal memory^116^. However, these segmentation effects were specifically facilitated by changes in emotional valence rather than arousal during sequence encoding, implying limited involvement of the LC. Because emotion was modulated by relatively pleasant musical pieces, it is possible that these boundaries lacked the intense spikes in arousal that may be necessary to engage the LC and impair temporal binding processes in memory. Differences between memory separation and integration could furthermore depend on whether a prediction error-related boundary is signed (i.e., positive or negative) or unsigned (i.e., absolute)^66^. Regarding generalizability, arousal processes have also been shown to track event boundaries in short stories, with greater boundary-related pupil responses predicting later memory for those narratives^117^. This suggests that pupil-linked arousal, and potentially LC activation, are sensitive to event structure across tasks of varying complexity. Our basic science findings will be enriched by future investigations using more real-world, naturalistic experiences, which may better capture the intricacies of everyday memory and neuromodulatory function.

## Lead Contact

Further information and requests for resources should be directed to and will be fulfilled by the lead contact, David Clewett.

## Materials Availability Statement

All stimuli are available on the first author’s Open Science Framework (osf.io/adbm4) page.

## Data Availability Statement

All behavioral and eye-tracking data are available on the first author’s Open Science Framework (osf.io/adbm4) page.

## Code Availability Statement

Code and scripts are provided on the first author’s OSF page (osf.io/adbm4).

## Acknowledgements

This project was funded by federal NIH grant R01 MH074692 to L.D. and by fellowships on federal NIH grant F32 MH114536 to D.C. We thank Ziyuan Chen for her assistance with locus coeruleus tracing. We thank Alexandra Cohen and Erin Morrow for helpful feedback on earlier versions of this manuscript. Additionally, we thank Nina Rouhani for stimulating conversations about these ideas. We dedicate this paper to Dr. Carolyn Harley, a treasured colleague and pioneer of locus coeruleus and memory research. The significant impact of her work is only outmatched by her generosity of spirit, kindness, and child-like enthusiasm for neuroscience research.

## Author Contributions

D.C. and L.D. conceptualized and designed the experiment; D.C. collected data; D.C., and R.H. analyzed the data; D.C and L.D. wrote the manuscript. R.H. helped revise the original manuscript.

## Declaration of Generative AI and AI-Assisted Technologies

The authors did not use AI tools in the preparation of this work.

## Supplemental Information

Document S1. Figures S1–S5

## STARS Methods

### Participants

Prior to the study, we performed a power analysis to estimate the appropriate sample size using the pooled temporal memory data from a very similar behavioral version of this event boundary experiment^16^. With an alpha = .05 and power = .80, we estimated that we would need 29 participants to obtain a large effect size (d = .80; Cohen’s criteria; G*Power 3.1).

Based on this estimate and to account for potential attrition, a total of 36 healthy young adults were recruited from the New York University (NYU) Psychology Subject Pool and nearby community to participate in this neuroimaging experiment. All participants provided written informed consent approved by the NYU Institutional Review Board and received monetary compensation for their participation. Eligibility criteria included having normal or normal-to-corrected vision and hearing, not taking beta-blockers or other psychoactive drugs, and having no bodily metal to ensure MRI safety.

### Data Exclusions

Of those 36 participants, four were excluded from all analyses due to falling asleep in the scanner (*n* = 3) or due to malfunction of the audio equipment (*n* = 1). This left a total of thirty-two participants (20 females; Mean_age_ = 22 years old, SD_age_ = 2.7 years) for all behavioral and fMRI analyses. Eleven participants reported being “White”, two reported being “Black/African American”, 15 reported being “Asian”, and 4 reported being “More than one race.” For the pupil-related analyses, four additional participants were excluded due to eye-tracker malfunction or poor eye-tracking quality, leaving a subset of twenty-eight participants with valid data for all brain, behavioral, and pupil measurements in this study. Sex differences were not analyzed or included as covariates in any analyses, because they were not central to our predictions and there were relatively small sample sizes of each type.

A small subset of participants did not complete all 10 blocks of the event sequence task because they chose to exit the scanner early: two participants completed 7 blocks, one participant had 8 blocks, and one participant had 9 blocks. All remaining data were usable and included in the analyses.

### Materials

The object stimuli consisted of 512 color images of everyday objects on a gray background. These images were selected from existing datasets^118,119^. Each image was resized to be 300 x 300 pixels for the encoding phase. For the temporal memory tests, the pair of test images were each resized to be 250 x 250 pixels to create more gaze separation on the screen. The luminance of all object images and fixation screens was normalized using the SHINE toolbox in MATLAB to control for non-cognitive-related effects on pupil size. A total of 320 images were used for encoding, 32 images were used for the practice block outside of the scanner, and 120 images were used as lures in the delayed item recognition test. The practice images were identical across all participants, whereas the encoding and lure items were randomized across participants.

For the auditory context manipulation during sequence encoding, six 1s pure tones with sine waveforms of different frequencies (500Hz, 600Hz, 700Hz, 800Hz, 900Hz, 1000Hz) were generated using Audacity (https://www.audacityteam.org). These frequencies were chosen because they were discriminable from one another and were arousing enough to maintain participants’ attention. They were also discriminable from the noise of the scanner.

### Overview of Protocol

This study involved one MRI session and one behavioral session ∼24 hours later. Upon arriving on Day 1, participants provided written informed consent and completed a demographics form. Next, participants were given instructions about the timeline of scanning and about the event sequence encoding task. They then performed one practice study-test block of the experimental task on a laptop.

### Scanning and behavioral procedures

Scanning took approximately 2.5 hours, involving the following sequences in order: one high-resolution anatomical scan, one T2-weighted scan, one LC MRI scan (a specialized sequence potentially sensitive to neuromelanin content), and 20 functional scans (10 study rounds interleaved with 10 temporal memory test rounds). Upon entering the MRI scanner, we calibrated the audio equipment while the T2 scan was performed. This audio test ensured that participants could discriminate between the different tone types and could hear the tones comfortably above the noise of the scanner.

Participants could adjust tone volume by providing button press feedback to the experimenter. Prior to each encoding list, participants were reminded of the button presses they should use to make their size judgements when viewing each image.

Participants returned to the lab approximately 24 hours later and performed a surprise item recognition memory test. This behavioral session lasted approximately 30 minutes. Given item recognition effects were not central to the hypotheses of the current study, those results are not reported in this manuscript.

### Event sequence encoding task

To determine if event boundaries shape the temporal structure of memory, we adapted a behavioral version of a novel paradigm that uses stability and change in auditory contexts to segment memories of neutral image sequences^16^ (**Figure 1A**). For each item sequence, participants viewed a series of 32 grayscale, luminance-normed images of objects. Each image was presented in the center of a gray background for 2.5 seconds. A black fixation cross was displayed in the middle of the screen in between each image for 3, 5, or 7 seconds. A 1-s pure tone was played half-way through each jittered ISI in participant’s left ear or right ear. This tone indicated to participants which hand they should use to judge if the object was larger or smaller than a standard shoebox (left ear = left hand). To promote associative encoding, participants were also encouraged to link sequential items together by creating a mental narrative.

To create a stable auditory context, or ‘event’, the specific tone/ear pairing heard before each object remained the same for eight successive objects. After the 8^th^ item in each auditory event, the tone switched to the other ear and changed in pitch, creating a theoretical ‘event boundary’ in the sequence. This new tone/ear pairing then remained the same for the next eight items, and so on. There were three auditory event boundaries per list, creating a total of four auditory events. Tone frequencies were pseudorandomized across lists such that no tones of a given frequency were presented more than once in a list (e.g., tones that were 700Hz were not heard in more than one event within a given list). Whether the tones first played in participants’ left or right ears was counterbalanced across lists. Additionally, 10 separate ISI orders were created, and the order of ISI sequence types across the task was randomized across participants. Each participant viewed a total of 10 lists/sequences in the scanner. Prior to entering the MRI scanner, participants performed one practice study-test block, which familiarized them with the task.

Notably, we specifically chose auditory cues because visual information, particularly luminance, inherently influence pupil size through the pupillary light reflex, which is a physiological response to changes in stimulus/screen luminance. The brightness of an image elicits changes pupil size unrelated to cognitive processes of interest. Further, the pupillary reflex is driven by activation of neuroanatomical pathways that are not regulated by the LC, including parasympathetic nervous system pathways that constrict the pupil^58^. Thus, using auditory cues enhanced our ability to isolate pupil changes that were not confounded by visual information (e.g., luminance), providing a more accurate, indirect readout of LC activation during the task.

### Delay distractor task

To create a 45-s study-test delay and reduce potential recency effects in memory, participants performed an arrow detection task after each sequence. In this phase, a rapid stream of either left-facing (<) or right-facing (>) arrow symbols appeared in the middle of the screen for 0.5s each. Each arrow was separated by a 0.5-s ISI screen with a central fixation cross. Participants simply had to indicate which direction the arrow was pointing via button press as quickly as possible.

### Temporal memory tests

Following the distractor task, participants performed two temporal memory tests. On each test trial, different pairs of items from the prior sequence were displayed on the screen for a fixed duration of 8s each. First, participants made a temporal order judgment by indicating which of the two items had appeared more recently during encoding (i.e., “which appeared later?”; **Figure 1**). Participants had four options based on the position of their choice on the screen, which were broken down by confidence: ‘definitely left’, ‘maybe left’, ‘maybe right’, or ‘definitely right’. Responses were made using separate button boxes placed in participants’ corresponding left and right hands. Participants then made a temporal distance rating in which they endorsed the item pair as having appeared ‘very close’, ‘close’, ‘far’ or ‘very far’ apart in the prior sequence (i.e., “how far apart?”). The two types of close responses were always made with the left button box and the two types of far responses were always made with the right button box.

Each temporal order and temporal distance test trial was also separated by a slower mini-version of the arrow distractor task, which provided an active baseline for fMRI analyses^120^. During inter-trial-intervals between test trials, a rapid stream of left-facing (<) or right-facing (>) arrow symbols appeared in the middle of the screen for 1s each. Each arrow was separated by a 1-s ISI screen with a central fixation cross. There could be 1, 2 or 3 arrows between each test pair, leading to a jittered temporal interval throughout the memory tests. Critically, each pair of items had always been presented with three intervening items during encoding. They were thereby always encountered the same objective distance apart. Because ISIs during encoding were jittered, we also pseudorandomized the timing of each to-be-tested pair window to always be 32.5s, such that the four ISI’s in this behaviorally relevant window summed to 20 seconds.

To test our hypothesis that event boundaries influence the temporal structure of memory, we examined two types of item pairs: (1) items that had appeared within the same auditory event (same-context pairs; 8 trials per list) and (2) items that had spanned an intervening tone switch (boundary-spanning pair; 6 trials per list). The list positions of the to-be-tested pairs were (B = boundary; SC = same-context): 1-5 (SC), 3-7 (SC), 6-10 (B), 8-12 (B), 9-13 (NB), 11-15 (NB), 14-18 (B), 16-20 (B), 17-21 (SC), 19-23 (SC), 22-26 (B), 24-28 (B), 25-29 (SC), and 27-31 (SC). The overall pair structure is displayed in **Figure S1.**

For all analyses, we removed the first tested item pair from encoding (positions 1 and 5), because it contained the first item in each list and likely constituted a task-irrelevant event boundary. Due to programming errors that were caught partway through the study, for a subset of participants, one block of the task was excluded from all analyses due to the ISI’s being 0.5s too short throughout the encoding sequence (*n* = 9 participants) and one boundary-spanning trial was excluded due to only having two rather than three intervening items (*n* = 23 participants).

### Logistic and linear mixed effects modeling analyses between brain and behavior

For all brain, behavioral, and brain-behavior correlation analyses, we performed linear and generalized linear mixed effects modeling analyses in RStudio (version 2022.07.1, R Core Team, 2017) using the *lme4* package^121^. Degrees of freedom and p-values were calculated using the *lmerTest* package^122^ The models were estimated using ML and Nelder-Mead optimizer. Random intercepts for Participant ID (Subject) were modeled as a random effect with a random intercept to control for individual differences. For the temporal order memory logistic regressions, judgments were collapsed across confidence ratings to increase statistical power. Order memory accuracy was then coded as a binary dependent variable (1 = correct, 0 = incorrect). Standardized parameters were obtained by fitting the model on a standardized version of the dataset. 95% Confidence Intervals (CIs) and p-values were computed using a Wald z-distribution approximation.

### Eye-Tracking Methods

#### Eye-tracking

Pupil diameter was measured continuously at 250 Hz during the event sequence task using an infrared EyeLink 1000 eye-tracker system (SR Research, Ontario, Canada). Raw pupil data, segmented by block, were preprocessed using ET-remove-artifacts, a publicly available Matlab program (https://github.com/EmotionCognitionLab/ET-remove-artifacts). This algorithm identifies blinks and other artifacts in the pupil timecourse, then either interpolates over these regions or imputes lengthy periods of artifacts with a missing data indicator (NaN).

Following the approach described in Mathôt et al. (2013)^123^, the algorithm detects blink events and other transient artifacts by identifying rapid changes in pupil size, or pupil velocity. The velocity timeseries is computed by applying MATLAB’s finite impulse response (FIR) differentiator filter on the raw pupil size timecourse, which provides a robust estimate of instantaneous rate of change while minimizing noise amplification. The parameters for the FIR filter (Filter Order = 14, Passband Frequency = 1, and Stopband Frequency=30) were chosen for this specific dataset (sampled at 250Hz) to ensure distinct and smooth trough-and-peak blink profiles in the velocity timeseries. MATLAB’s findpeaks function is then used to identify peaks and troughs in the pupil velocity timecourse. The Peak and Trough Threshold Factor and Trough Threshold Factor, which sets the minimum height constraint to qualify as a peak, were set at 3 or 4 standard deviations of the velocity timeseries, depending on the frequency of blinks for each subject. A contiguous trough followed by a peak in the velocity timeseries was identified as a blink profile.

For artifact removal, linear interpolation is applied across identified blink intervals. Artifact intervals greater than 2 seconds were automatically imputed with NaN (missing data indicator). After applying the algorithm with these settings, a trained user (R.H.) qualitatively inspected the output and occasionally used the ET-Remove-Artifact’s Manual Edit functionality. During this process, lengthy periods (>1s) of noisy pupil data were imputed with NaN, and sharp spikes or troughs (<1s in width) missed by the algorithm were interpolated over. A by-participant boxplot outlier approach was used to remove outlier trials from the dataset.

#### Average boundary-induced pupil dilation analysis

To determine if event boundaries elicit a transient increase in pupil-linked arousal, we compared tone-evoked pupil responses to the boundary tone (i.e., tone switch after the 8^th^ item in an event) and the same-context, or repeated, tones (i.e., tones that repeated before items 2-8 in a stable auditory event). Tone-evoked pupil dilation was computed as the average pupil diameter 1-1.5s after tone onset minus the average pupil size during the 500ms window prior to tone onset (**Figure 4A**).

This time window was chosen because it captured the same post-tone fixation screen shared across all trials and was not confounded by the onset of the ensuing object image. We then performed a linear mixed modeling analysis to test for differences in pupil dilation elicited by boundary versus same-context tones.

#### Pupil dilation temporal principal component analysis (PCA)

A temporal PCA was used to dissociate distinct autonomic and functional components of stimulus-evoked pupil dilations^16^. For each participant and type (boundary tones and same-context tones), we computed the average time-course of baseline-normed pupil dilations across all 1.5-s post-tone time windows. This resulted in 56 input variables (28 participants with one input per condition) to the PCA that contained 375 pupil samples each (see **Figure 4C)**.

An unrestricted PCA using the covariance matrix with Varimax rotation and Kaiser normalization was used to generate meaningful pupil components and component loading scores. These pupil component loadings index temporally dynamic, correlated patterns of pupil dilation elicited by the tones. Factor loadings with eigenvalues greater than 1 were retained and analyzed in subsequent analyses (Kaiser criterion^124^). These PCA loadings reflect the relative degree of engagement of that specific feature of pupil dilation. To determine if boundaries modulated different temporal characteristics of pupil dilation, we performed two-tailed paired t-tests on the loading scores for each pupil component with an alpha = .05 (**Figure 4D**).

### fMRI Acquisition and Preprocessing

#### fMRI/MRI data acquisition

All neuroimaging data were acquired with 3T Siemens Magnetom PRISMA scanner using a 64-channel matrix head coil. Scanning commenced with a high-resolution MPRAGE T1-weighted anatomical scan (slices = 240 sagittal; TR = 2300ms; TE = 2.32 ms; TI = 900 ms; FOV = 230 mm; voxel in-plane resolution = 0.9 mm^2^; slice thickness = 0.9 mm; flip angle = 6°; bandwidth = 200 Hz/Px; GRAPPA with acceleration factor = 2; scan duration: 5 min. and 21 s).

This scan was then followed by a T2-weighted scan (slices = 240 sagittal; TR = 3200ms; TE = 564 ms; FOV = 230 mm; voxel in-plane resolution = 0.9 mm^2^; slice thickness = 0.9 mm; flip angle = 6°; bandwidth = 200 Hz/Px; GRAPPA with acceleration factor = 2; scan duration: 3 min. and 7 s). During this scan, we also tested and calibrated the audio equipment to ensure the participant could hear the task-related tones above the noise of the scanner. A pair of fieldmap scans were also acquired to aid with functional imaging unwarping, with one scan acquired in the AP phase encoding direction and the other in the PA phase encoding direction. Prior to the encoding task, we collected a T1-weighted fast spin echo (FSE) imaging sequence to image LC structure (TR = 750 ms; TE = 12ms, voxel in-plane resolution = 0.429 × 0.429 mm^2^, slice thickness = 2.5 mm, slice gap = 3.5 mm; flip angle = 120°, 11 axial slices, FOV = 220 mm, bandwidth = 220 Hz/Px).

Separate functional images were collected for each of the interleaved 10 encoding runs and10 retrieval runs of the task. These images were acquired using a single whole-brain T2*-weighted multiband echo planar imaging (EPI) sequence (128 volumes per encoding run; TR = 2000ms; TE = 28.6 ms, voxel in-plane resolution = 1.5 x 1.5 mm^2^; slice thickness = 2 mm with no gap; flip angle = 75°, FOV = 204mm X 204mm; 136 X 136 matrix; phase encoding direction: anterior-posterior; GRAPPA factor = 2; multiband acceleration factor = 2). In each volume, 58 slices were tilted minus 20° of the AC-PC and were collected in an interleaved order.

#### fMRI preprocessing

Image preprocessing was performed using FSL Version 6.00 (FMRIB’s Software Library, www.fmrib.ox.ac.uk/fsl). Functional images were preprocessed using the following steps: removal of non-brain tissue using BET, B0 unwarping using fieldmap images, grand-mean intensity normalization of the 4D data set by a single multiplicative factor, and application of a high-pass temporal filter of 100s. No spatial smoothing was applied to preserve the spatial specificity of anatomical ROI’s and to improve pattern similarity estimates^125^.

Motion correction was performed in multiple steps. First, head movements were first detected using the FSL MCFLIRT tool, resulting in six motion nuisance regressors. Second, we computed DVARS (D referring to the temporal derivative of the time series, VARS referring to the root-mean-square variance over voxels) using the fsl_motion_outliers tool. DVARS quantifies the framewise change in BOLD signal intensity across the brain, providing a measure of temporal signal variability. Outlier volumes were detected using a boxplot method, where a volume was flagged if its DVARS value exceeded 1.5 times the interquartile range (IQR) above the third quartile. The labeled outlier volumes were subsequently included as nuisance regressors in the GLM’s to reduce their influence on the statistical analyses. Third, entire fMRI blocks with an average framewise displacement > 0.75mm, or half the voxel size, were excluded from analysis (across entire dataset = 12 runs). This block-level exclusion criterion was used to estimate general levels of restlessness or fatigue across an entire item sequence.

Each participant’s denoised mean functional volumes were co-registered to their T1-weighted high-resolution anatomical image using brain-based registration (BBR). Anatomical images were then co-registered to the 2mm isotropic MNI-152 standard-space brain using an affine registration with 12 degrees of freedom.

Due to its small size and location next to the fourth ventricle, BOLD signal in the LC is especially susceptible to physiological artifacts. To address this, eight separate physiological nuisance signal regressors were extracted for the subsequent GLM analyses. First, FSL FAST was used to decompose each participant’s high-resolution anatomical images into probabilistic tissue masks for white matter (WM), grey matter (GM), and cerebrospinal fluid (CSF). The CSF and WM masks were thresholded at 75% tissue-type probability to increase their spatial specificity and reduce potential overlap. Following a similar approach to Barton et al. (2019)^126^, we defined eight 4-mm spheres in representative regions of WM and CSF (four of each type; for exact coordinates, see Barton et al., 2019). Importantly, one of these spheres included a location in the fourth ventricle located adjacent to the locus coeruleus, which helped us mitigate artifact related to cardiac and brainstem pulsation in fMRI scans. The eight spheres and WM and CSF anatomical masks were then transformed into each participant’s/run’s native functional space and merged to increase their spatial specificity even further.

Nuisance timeseries for each of the four WM and four CSF merged masks were then extracted from each run’s preprocessed functional data.

### Locus Coeruleus and Hippocampal Subfield Region-of-Interest (ROI) Definitions

#### Locus coeruleus ROI

To acquire participant-specific anatomical LC masks, we collected specialized structural scans that are thought to be related to neuromelanin concentration. LC neurons contain neuromelanin, a byproduct of NE metabolism, thereby enabling its localization via specialized imaging sequences (see **Figure 5D** for example participant’s LC MRI scan). To localize and delineate the LC in each participant, LC ROIs were hand-drawn on each participant’s LC MRI scan using a similar procedure to a previous study^127^. Bilateral LC anatomical ROIs were manually defined as a three ∼1.29 mm wide by ∼1.29 mm long masks in three adjacent axial slices where LC signal was brightest and most visible. To identify the most superior slice with the LC, we transformed a consensus anatomical LC mask from an existing dataset into each participants’ native LC MRI image space^128^. The most superior axial slice was defined as highest axial slice that contained this anatomical reference ROI of the LC. For all three axial slices, the LC ROI masks were centered upon the left and right brainstem voxels with the highest MR signal intensities neighboring the corners of the fourth ventricle. We also drew a separate reference mask for the dorsal pontine tegmentum (PT) in each of the three slices, which would later be used to account for overall noise across the images in the neuromelanin signal intensity calculations. This PT reference anatomical ROI was defined as a 10 × 10 voxel square located 6 voxels above the more ventral of the 2 LCs and equidistantly between them.

All masks were hand-drawn by two individuals trained on the anatomy of the LC (R.H. and Z.C.). Their drawings showed high inter-rater reliability (ICC = 0.97). Average LC contrast-to-noise ratio (CNR) was 0.17 and the standard deviation was 0.027, consistent with previously reported CNR values using this fast spin echo MRI sequence (e.g.,^101,127^). LC CNR was not significantly correlated with age (ρ = 0.11, p = 0.56) nor did it significantly differ by sex (t(30) = -1.65, p = .11, Cohen’s d = -0.60).

For the fMRI and ROI analyses, each participant’s LC MRI image was first brain-extracted using BET and the small field of view parameter (-Z). These brain-extracted LC MRI scans were then co-registered to each participant’s high-resolution anatomical scan using an affine transformation with 6 DOF. We then performed two separate registrations using the inter-space transformation matrices acquired during image preprocessing.

Namely, the hand-drawn LC ROIs were first transformed from native space to anatomical space and then from anatomical space to each participant’s run-specific native functional space. All fMRI analyses of the LC were conducted in native functional space.

#### Hippocampal subfield segmentation and quality control

Bilateral hippocampal subfields CA2/3, DG, and CA1 were segmented from each participant’s high-resolution anatomical scan using Freesurfer 6.0 (https://surfer.nmr.mgh.harvard.edu/). The T2-weighted images were also used to facilitate segmentation. The Freesurfer 6.0 hippocampal segmentation uses Bayesian inference based on observed image intensities and a probabilistic atlas built from a library of in vivo manual segmentations and ultra-high resolution (∼0.1 mm isotropic) ex vivo labeled MRI data^113,129^. A multi-step quality control procedure was used to ensure accuracy of the subfield segmentations. We first used automated measures computed by Freesurfer of the Euler number (a metric of cortical surface reconstruction) to classify poor quality structural scans^130,131^. We excluded outliers who on a box-and-whisker plot were above Q3 + 3 * the interquartile range. This step resulted in no exclusions.

We next followed the proposed quality control procedure guidelines for FreeSurfer-based segmentation of the hippocampal subregions designed for the Enhancing Neuro Imaging Genetics through Meta-Analysis (ENIGMA) consortium^114^. We first checked for outliers (+/− 2 SDs) for each hippocampal subfield volume, total brain volume, total GM volume, ICV, and GM/ICV ratio^115^. To be as conservative as possible, participants that had outliers in any of these measures was excluded from all hippocampal analyses. This process identified the following four hippocampal subfield volume outliers (all separate participants): left CA1 for one participant, left CA3 for one participant, right DG for one participant, and right CA2/3 for two participants. Furthermore, three outliers were also flagged for the GM/ICV ratio: one of those participants overlapped with outliers identified in the hippocampal subfield checks, and two additional new participants were flagged as outliers in this specific step. These outlier detection steps resulted in the removal of 7 participants from all hippocampal analyses, resulting in N = 25.

Next, we checked the processed data quality using the rank-order rules of subfield volumes, and also plotted the histograms of each measure^114^. As a final step, we conducted HTML-snapshot-based visual QC of hippocampus segmentation of all participants, with particular attention paid to the hippocampus mask as a whole and hippocampus fissure within the hippocampal mask. For visual inspection, we followed QC guidelines from Yoo et al. (2023)^115^ and Sämann et al., (2020)^114^: 1) Is binary hippocampal mask visible? 2) Is hippocampal fissure positioned within hippocampal mask? 3) Are larger portions of the hippocampus cut off? 4) Are there any subfield-related peculiarities? There was no additional exclusion after this QC pipeline, as the only problematic participants overlapped with the original subfield volume checks (see above). We did not conduct any manual edits to avoid introducing external bias or human error.

Validated hippocampal ROIs were co-registered to each participant’s native/run-specific functional space for subsequent fMRI pattern similarity analyses. These native-space hippocampal masks were then thresholded at 0.2 to reduce spatial overlap between adjacent subfields. Each subfield was separated into left and right hemisphere masks based on evidence of potential lateralization of memory processes in hippocampus.

### FMRI Analyses

#### Generalized linear modeling (GLM) analyses and acquisition of single-trial beta estimates of brain activation during the encoding task

One of the main goals of this study was to test if event boundaries alter responses in the LC and hippocampus, and whether such engagement relates to changes in arousal systems and how individuals remember the order and timing of recent events. To this end, we first performed Least Squares Separate (LSS) GLM analyses to acquire single-trial estimates of brain activation, or beta maps, across the whole brain^132,133^. These GLM’s were performed on unsmoothed functional data and in each participant’s native functional space for each encoding run, separately.

In the LSS procedure, each tone and image from a given sequence was modeled as its own trial-of-interest in separate GLM’s, resulting in a unique beta map for each stimulus (n = 64 stimuli per list; 32 tone trials and 32 image trials). Within each single trial GLM, tones were modeled as a stick function with a duration of 1s, while each object image was modeled as a stick function with a duration of 2.5s. The first regressor in each GLM represented the trial of interest (*n* = 1), while the second regressor modeled all other trials (*n* = 63). This modeling process was performed iteratively to generate unique beta maps for each image and tone in the encoding lists. To account for movement artifacts and physiological noise, each GLM included 14 nuisance regressors (4 WM regressors, 4 CSF regressors, and 6 motion regressors) as well as additional regressors for individual volumes that were flagged for extreme head movements.

Next, we extracted trial-level betas (i.e., parameter estimates) for each tone trial from each participant’s LC anatomical mask. Single-trial beta estimates are often noisy due to the transient effects of head motion, cardiac pulsation, and other MRI-related artifacts^125^. To account for potentially spurious estimates of brainstem activation, outlier trials were identified at the participant level using a boxplot outlier removal method. This resulted in the removal of 1.75% of the tone-evoked LC trials from the entire dataset.

#### fMRI analyses of average tone-evoked LC activation during encoding

To test if event boundary tones elicited LC activation, we performed linear mixed modeling analyses using the lmer4 package in R. For the logistic regression analysis on temporal order memory, Condition was modeled as a fixed-effect predictor of LC activation (boundary-spanning = 1; same-context = -1). The outlier-cleaned LC responses were mean-centered by participant and then entered as fixed-effects predictors of temporal order memory. Order memory was coded as a binary outcome variable (1 = correct and 0 = incorrect). Subject ID and the side of the screen with the correct answer were modeled as random effects with a random intercepts and constant slopes.

#### Hippocampal pattern similarity analyses

To test if event boundaries reduce the stability, or similarity, of multivoxel hippocampal representations across time, we performed a multivariate pattern similarity analysis. For each of the six hippocampal subfield ROIs, we first extracted activation patterns from the trial-unique beta maps produced by the LSS GLM (see **Figure 3A** for schematic; also see **Figure S1** for a schematic of to-be-tested pair structure). Pattern similarity scores were then computed at the item pair level by correlating multivoxel patterns between each of the to-be-tested trial pairs from encoding. For example, we extracted the average multivoxel pattern for images in position 3 and position 7 in each list and then correlated these patterns. This Pearson correlation, or pattern similarity (PS), score provided a neural measure of how similar hippocampal subfield activity patterns were across encoding. As such, lower hippocampal PS values index greater temporal pattern separation, whereas larger PS values index greater pattern integration.

Importantly, the spacing between to-be-tested memory item pairs was large (32.5 seconds) and exceeded the time course of the canonical hemodynamic response function (HRF). This time window was always identical across all pair types, thereby mitigating potential issues of temporal autocorrelation in the BOLD signal. As before, we used a by-participant boxplot outlier removal method to exclude spurious or noisy PS trials from analysis^125^. This outlier filtering method had a minimal impact on data exclusions, only removing a range of 0.90-1.56% of the PS datapoints across the 6 subfields. The remaining hippocampal PS scores were mean-centered by participant and modeled simulatenously as fixed-effects predictors of Condition (1 = boundary, 0 = same-context). Subject modeled as a random effect with a random intercept. To examine if hippocampal pattern stability related to temporal order memory, we performed the same multiple logistic mixed effects modeling analyses as for the LC, with all six subfields modeled as fixed effects.

To examine if hippocampal pattern stability relates to transient activation of the LC at event boundaries, we performed multiple linear mixed effects models. Here, tone-evoked LC activation, or parameter estimate, was modeled as a continuous outcome variable. Pairwise pattern similarity values for all six subfields were mean-centered by participant and modeled as fixed-effects predictors of LC activation. Condition and its interaction with each subfield’s PS values were also entered as fixed-effect predictors. Subject ID was modeled as a random effect with a random intercept (see **Figure 3A** for schematic of analyses). The alignment between timepoints of tone-evoked LC activation and their corresponding hippocampal PS item pairs is displayed in **Figure S1.**

#### fMRI analyses of low frequency fluctuations in LC signal and both LC structure and online arousal processes

Even though fMRI cannot be used to measure absolute levels of activity, researchers have endeavored to find other indirect measures of tonic LC activity. Hypothetically, tonic LC activity reflects slower and largely task-unrelated patterns of brain activity, arousal, and attention^33,62^. Thus, measuring LC blood-oxygen-level-dependent (BOLD) signal fluctuations after removing task-related patterns may provide an indirect index of tonic LC activity. Inspired by this idea, we performed additional exploratory analysis to examine if ostensible patterns of tonic LC activation were meaningfully related to LC contrast-to-noise ratio (CNR) as well as our other pupil, brain, and behavioral measures. We note that these methods are very exploratory, and primarily follow the logic that tonic LC activation is most likely indexed by low-frequency, task-unrelated fluctuations in BOLD signal across the task.

Past fMRI work has examined how low frequency fluctuations in the LC BOLD signal is functionally coupled with activation patterns in other brain regions while participants rest. It is thought that these patterns could index “tonic” patterns of LC functional connectivity due to the absence of meaningful, task-related information^51,134^. One recent fMRI study measured low frequency BOLD variance (LFBV) during resting-state fMRI in key nodes of the ventral attention network (VAN) as a measure of cortical reactivity related to LC tonic activity^63^. They then related this indirect measure of tonic LC activity to online measures of task-evoked pupil dilations during a digit span task. They found that, across participants, higher tonic low frequency BOLD variability in the VAN at rest corresponded with reduced task-induced pupil dilations^63^. In sum, this finding is highly consistent with the idea that tonic levels of LC activation constrain the magnitude of phasic, task-related LC responses. In this case, highly elevated levels of tonic LC activation should reduce LC phasic responses, as evidenced by smaller pupil dilations. It is possible that low-frequency BOLD signal fluctuations help capture this tonic LC effect. For instance, work in rodents shows that restraint-induced stress, which should increase background arousal levels, can augment infraslow fluctuations in LC activity (0-0.5Hz range^86^).

Following this logic and analytical approach, we performed two different exploratory analyses of tonic LC activation. Based on prior work, we extracted low-frequency BOLD variability from the LC after regressing out task-related patterns of brain activation. Using the functional data from the auditory sequence encoding task, we first created separate event-related regressors by modeling the onset times of all tones and images with durations of 1s and 2.5s, respectively. These task regressors were also separate into boundary trials or same-context trials. Each task regressor was convolved with a dual-gamma canonical hemodynamic response function and their temporal derivatives were used to model the data in a whole-brain GLM. We also modeled the same noise regressors as our LSS-GLM analyses, which included nuisance regressors for motion, extreme head movements, and both WM and CSF signals.

Next, we extracted the residuals from these GLM analyses, which preserved fluctuations in the BOLD signal that were not tied to the stimuli or physiological noise. These GLM’s were performed separately for each participant and for each individual block of the encoding task. Each of the residual brain maps output by the GLM’s was then band-pass filtered into the 0.01-0.1Hz range to isolate low-frequency components of the BOLD signal. To estimate patterns of tonic LC activation, we performed an ROI analysis on these brain maps by extracting low frequency BOLD signal from each participant’s LC anatomical mask (for a sample timeseries, see **Figure S5A**). To further ensure that these block-level measures of LC BOLD activation were not driven by noisy head movements in general, we excluded data from any blocks where the absolute framewise displacement values exceed 0.75mm.

Tonic LC activity is related to sustained, baseline arousal states, as opposed to phasic activity, which involves brief, task-related bursts of activity^62^. We therefore reasoned that higher variability in the low frequency range may capture tonic LC activation, where the LC is influencing ongoing arousal levels over a longer period. Here, we assessed tonic LC activation in two different ways. First, we computed the standard deviation of these low-frequency BOLD signals across each task block, providing an indirect measure of the level of slower, task-unrelated engagement of the LC^63^. We term this measure “low frequency (LF) bold variability”.

Second, we computed the power of lower-frequency BOLD signal in the 0.01-0.1Hz range as an estimate of the level of tonic LC BOLD activation across the task. The time series for each voxel was transformed to the frequency domain and the power spectrum was obtained. Since the power of a given frequency is proportional to the square of the amplitude of this frequency component, the square root was calculated at each frequency of the power spectrum and the averaged square root was obtained across 0.01–0.1Hz at each voxel. This averaged square root was taken as the amplitude of low-frequency fluctuations, or “ALFF”^135^. The global mean of ALFF was then extracted across all voxels using a binarized whole-brain mask for each participant. The local ALFF values from the LC signal was then standardized by dividing by the global ALFF mean, providing a final standardized index of LC tonic activation.

Having now acquired these two functional measures of BOLD fluctuations in the LC, our primary goal was to cross-validate different indirect measures of LC tonic activation with each other. To this end, we performed Spearman’s rho partial correlation analyses between our two functional measures of tonic LC activation, LF BOLD Variability and ALFF, and our other key pupil and LC measures. Significance was assessed at a one-tailed p-value < .05 due to our strong a priori predictions that LC tonic activation would be anti-correlated with phasic measures of LC activation and pupil dilation. All results from these individual difference correlation analyses are displayed in **Figure S5**.

### Individual differences correlation analyses

Individual differences analyses were performed between measures of temporal memory (order and distance), pupil-linked arousal, pupil dilation component loadings, encoding response times, LC activation, and LC contrast-to-noise ratio (CNR) using Spearman’s rank order correlations (see **Figure 5A**). We were specifically interested in isolating boundary-evoked effects on all these neurophysiological and behavioral variables. Thus, for each participant, we computed difference scores by subtracting the average values across all same-context trials from the average values for boundary trials for encoding RT’s, pupil dilation loadings from the PCA, and tone-evoked LC activation. We also computed difference scores for order accuracy by subtracting average performance for all same-context pairs from average values for all boundary-spanning pairs. All these measures were furthermore correlated with LC CNR scores to examine their relationship with LC structural integrity. For exploratory correlations with our two “tonic” LC activation measures, we performed separate Spearman’s rank order correlations with one-tailed p-values due to the strong directional predictions of these analyses (see **Figure S5**).

---

**Supplementary Figure 1.**
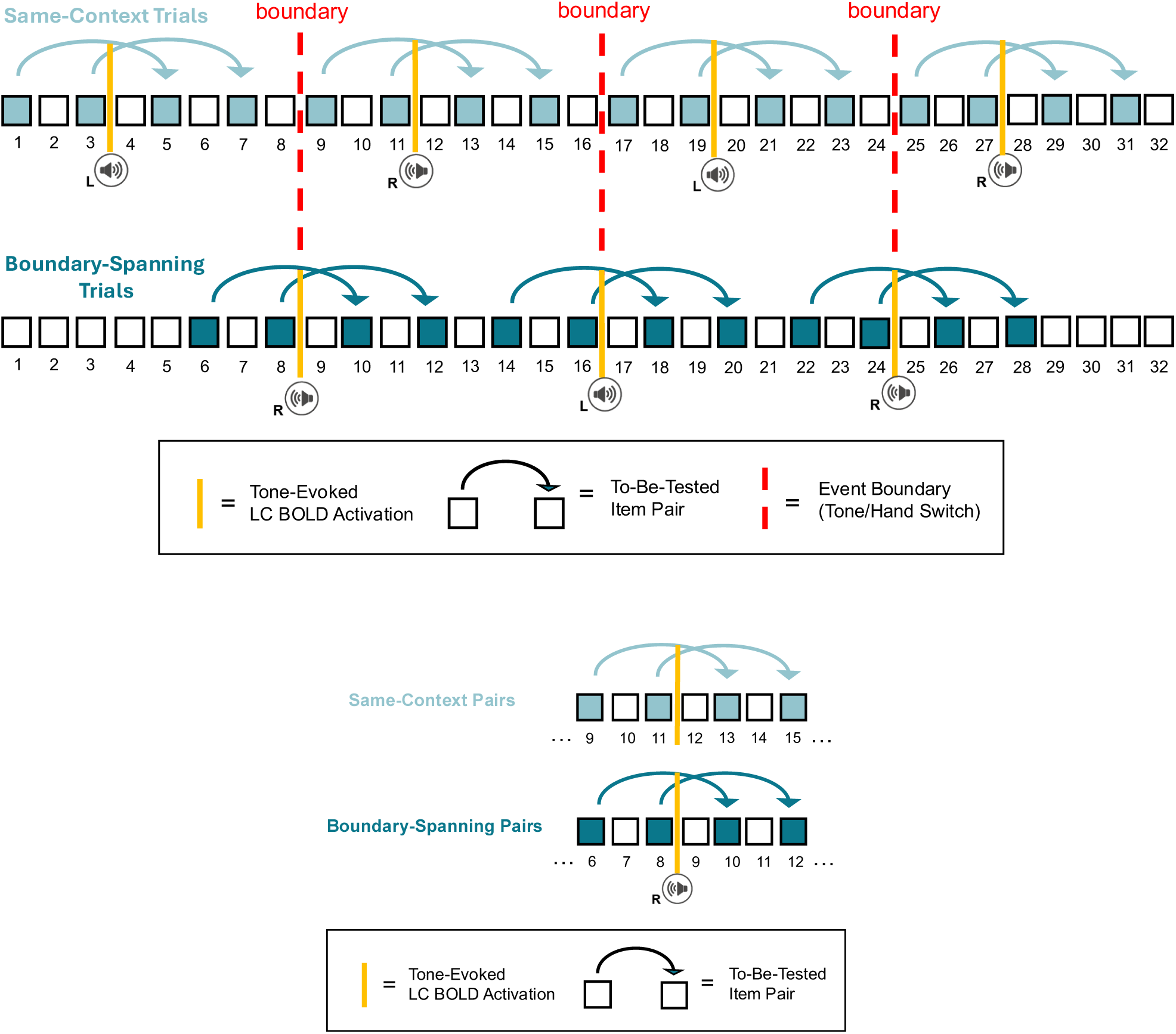
Auditory event sequence structure and analysis schematic, related to Figure 1 (Top Panel) Each sequence contained 32 images of everyday objects. A pure tone was played either in participants’ left ear or right ear for 8 successive items. It then switched to the other ear and changed in pitch. The side/tone then repeated for another 8 items before switching ears again and so on. Light blue colored squares denote the item pair positions that were subsequently queried during the temporal memory tests after each list. The arrow connections indicate which item pairs were later tested together, in the memory test. There were always three intervening items between each to-be-tested pair during encoding. Dark blue squares show the pair positions for boundary trials, or to-be-tested item pairs that spanned an intervening tone switch. Vertical dashed red lines indicate the positions of “event boundaries”, or the three auditory tone switches in each list. For the linear and logistic mixed effects modeling regression analyses, we aimed to relate trial-level estimates of tone-evoked LC activation to both temporal order memory performance and hippocampal pattern similarity for their corresponding item pairs. To align these two measures appropriately, we specifically focused on LC parameter estimates (derived from BOLD signal model fits in the GLM) evoked by event boundaries and same-context tones positioned-matched to those locations (all vertical gold lines). (Bottom Panel) Example of position-matching between tone-evoked LC activation between conditions. This illustration shows how the specific tones that were analyzed for LC activation were position-matched based on the location of event boundaries. This provided the critical reference point for controlling timing-related effects of LC activation relative to the positioning of the to-be-tested item pairs spanning those moments. In this example section from an item sequence, the first boundary tone occurred after the 8^th^ item in the list. This means that the boundary occurred after two items for the item from the first pair that spanned those boundaries (i.e., item in position 6), and immediately after the first item in the second boundary-spanning item pair (i.e., item in position 8). To position-match these timing effects in the same-context condition, the two same-context pairs were also temporally aligned with the same sampling points. That is, the relative positioning of the selected tones in the analysis was the same as the two boundary-spanning pairs for each event boundary in the list.

**Supplementary Figure 2.**
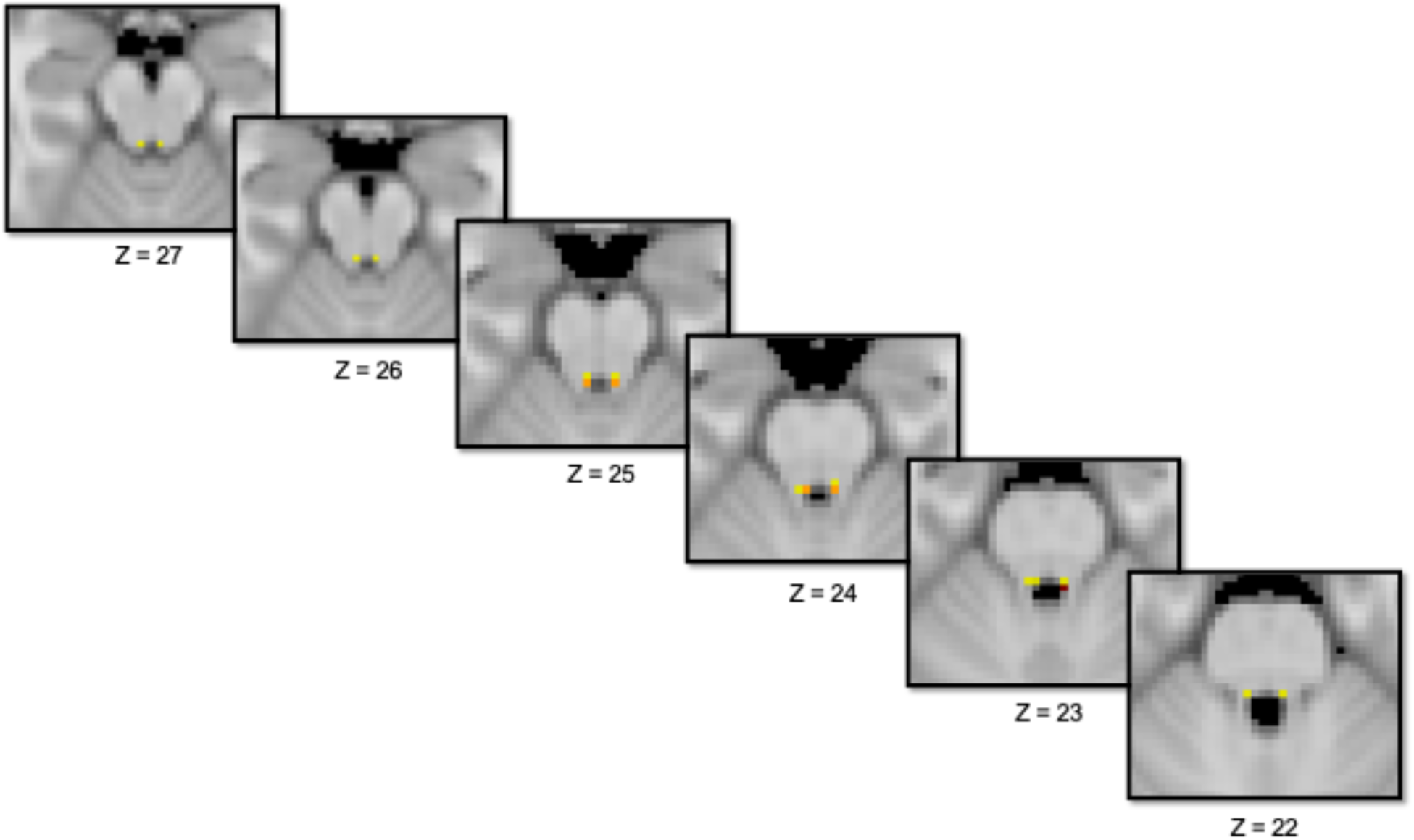
High spatial correspondence between a consensus locus coeruleus (LC) mask constructed from our LC MRI images and a standard LC mask from the literature, related to Figure 2. Yellow voxels = LC mask from Dahl et al. (2022)^1^; red voxels = consensus mask from current study; orange voxels = spatial overlap between voxels in both masks. Z values indicate slice location in standard space in the axial plane.

**Supplementary Figure 3.**
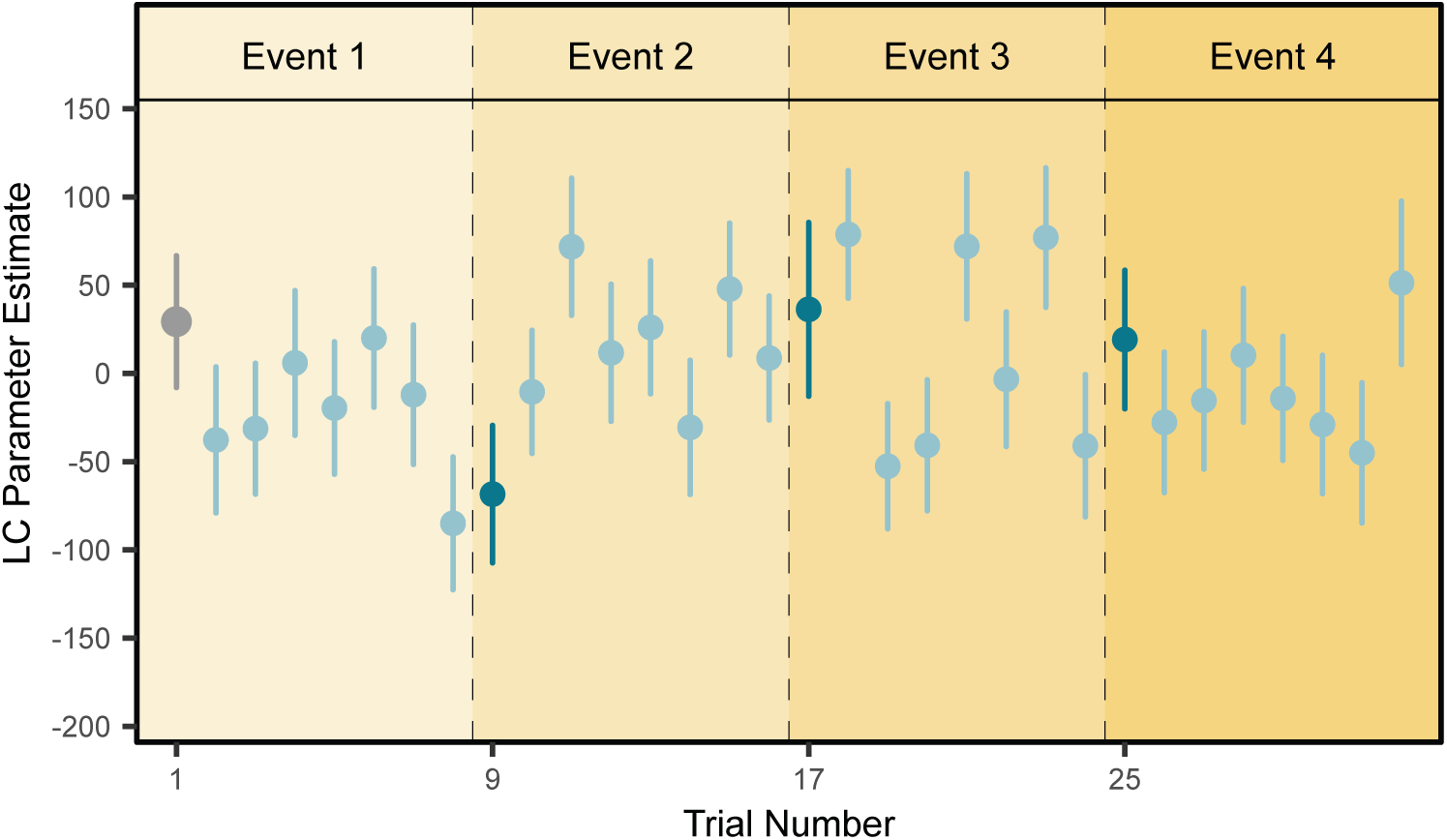
Patterns of locus coeruleus (LC) activation across the thirty-two tones in the sequences, related to. Figure 2. Average LC parameter estimates, or estimates of activation levels, across all the tones, participants, and lists. The first trial in the list represents a task-irrelevant event boundary and is displayed in grey. Dark blue colors in positions 9, 17, and 26 signify the positions of the three tone switches, or event boundaries, in each list. The lighter blue colors represent the repeated, or same-context tones, that helped define each auditory event (27 repeated tones in total, excluding the first item in list). Dots represent means and bars represent s.e.m. The data includes n = 32 participants.

**Supplementary Figure 4.**
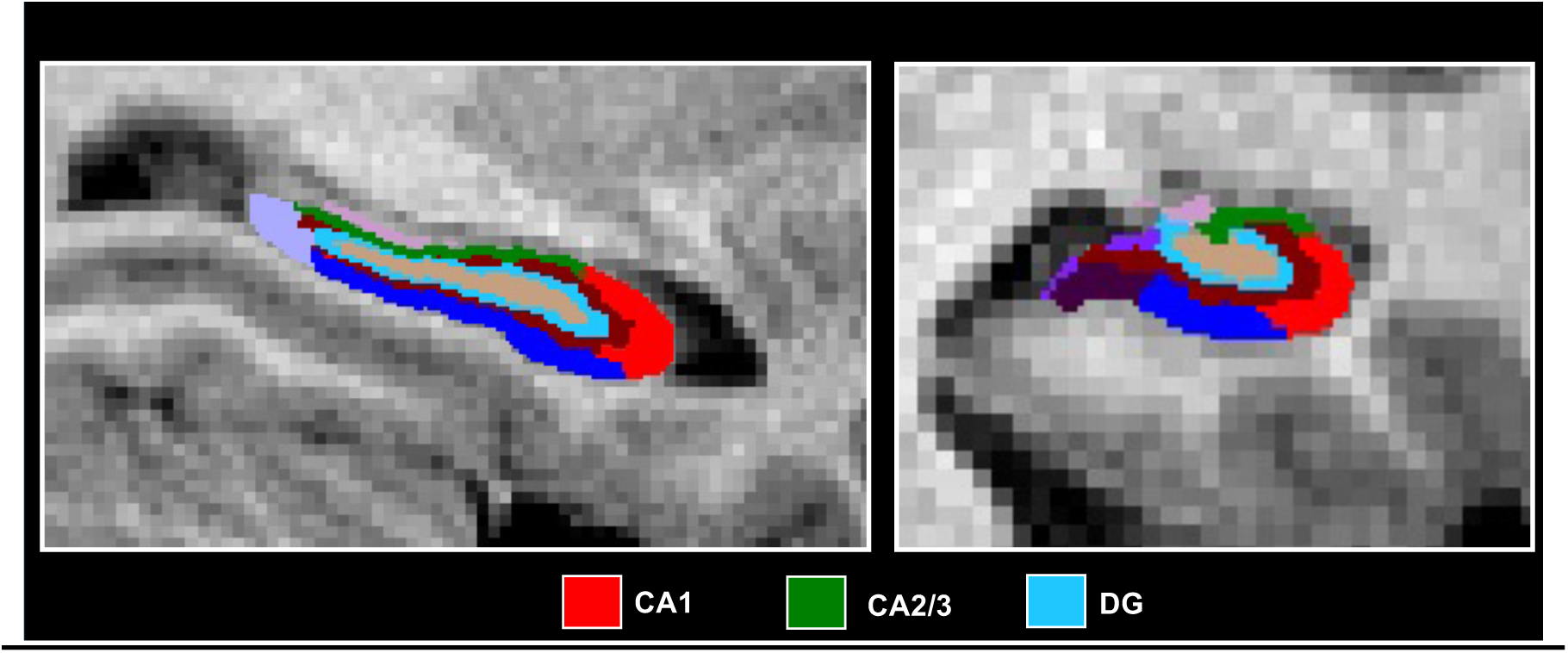
Example participant’s high-resolution T1 anatomical hippocampal subfield segmentation from Freesurfer 6.0, related to Figure 3. The left panel shows a sagittal slice of the segmentation along the long axis of the hippocampus. The right panel shows a sample coronal slice. Turquoise = dentate gyrus (DG); red = CA1; green = CA2/3.

**Supplementary Figure 5.**
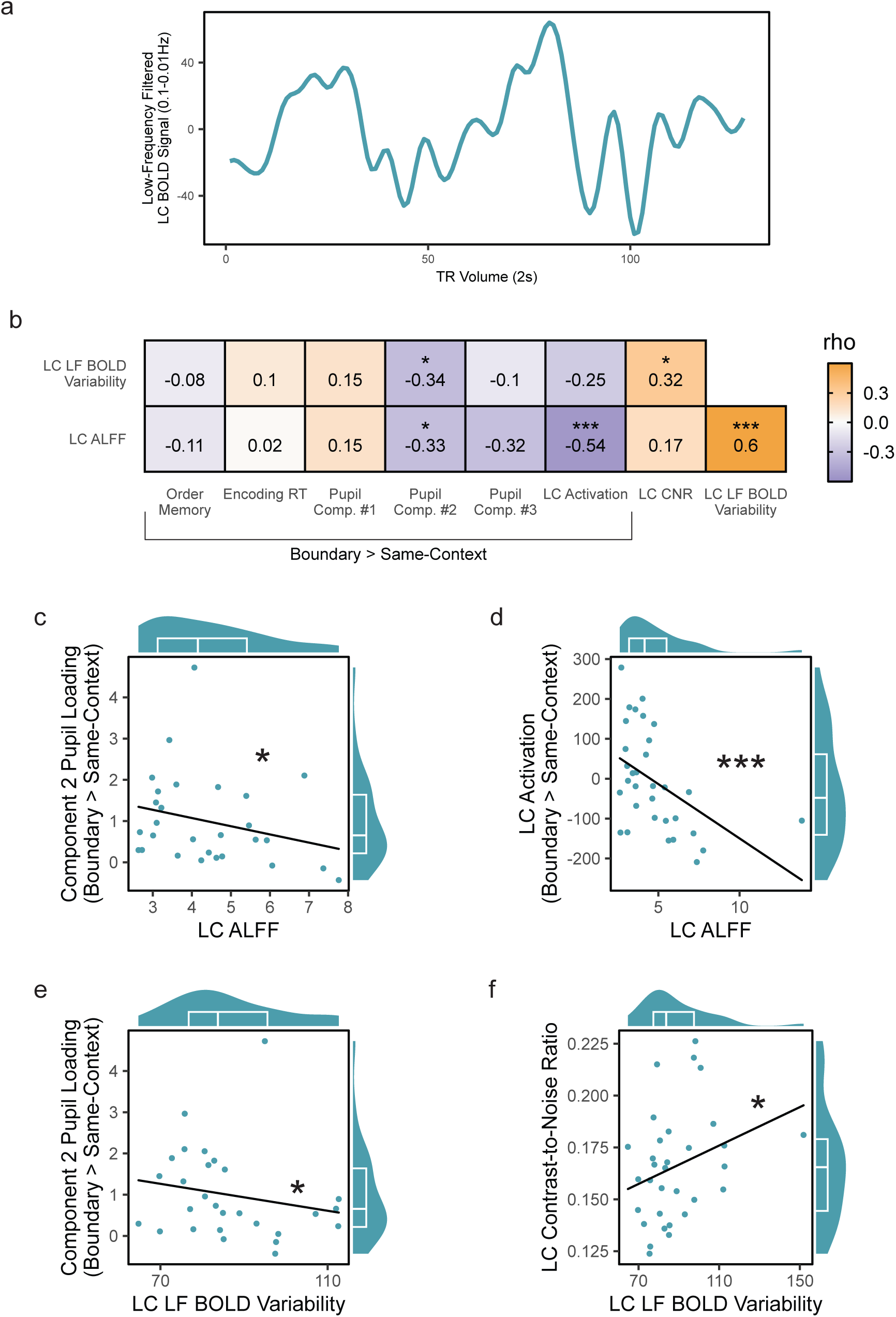
Structural and functional measures of elevated background, or tonic, LC activation are anti-correlated with phasic measures of boundary-induced arousal and LC responses, related to Figure 5. (A) An example subject’s timeseries of low-frequency BOLD fluctuations in the LC after regressing out task-related brain activity and bandpass filtering the residual signals between 0.01-0.1Hz. (B) Spearman’s rank order cross-correlation matrix relating two low-frequency BOLD signal fluctuation measures to other key functional, pupil dilation, and behavioral variables. These two BOLD measures included “low frequency (LF) BOLD Variability,” or the standard deviation of the BOLD fluctuations across the timeseries, and “analysis of low frequency fluctuations”, or ALFF, which measures the power of the LC signals in this frequency domain. For all functional and behavioral measures, we used subtraction scores of boundary trials versus same-context trials to isolate boundary-specific effects. Pupil variables included loadings from the three aspects of pupil dilation identified by the temporal PCA. Task-related, phasic noradrenergic activity was assessed using tone-evoked changes in LC BOLD activation. Behavioral metrics of memory separation included response-time (RT) slowing at boundaries and temporal order memory impairments. LC structure was assessed by extracting the contrast-to-noise ratio (CNR) in each participant’s fast spin echo MRI images. Within the correlation matrix, colored bar indicates the Spearman’s rho correlation coefficient with purple values indexing negative correlation coefficients and orange boxes indexing positive correlation coefficients. Color saturation reflects the strength of the correlations. (C/D) Spearman’s rho correlation plots showing that higher LC ALFF was significantly anti-correlated with boundary-induced loadings on pupil component #2 (C) and higher boundary-induced LC activation (D). (F) Spearman’s rho correlation plots showing that higher LC LF BOLD Variability was positively correlated with higher LC CNR and anti-correlated with boundary-induced loadings on pupil component #2. For all correlation plots (panels C-F), marginal density-boxplots depict the median (center line) and interquartile range (25th–75th percentiles) of the x- and y-distributions, and individual dots represent each participants’ data. All behavioral and brain correlations have n = 32 participants; any pupil-related correlations have n = 28 participants. Significance is indicated with a one-tailed test. *p < .05, ***p < .001.

